# WaveSeekerNet: Accurate Prediction of Influenza A Virus Subtypes and Host Source Using Attention-Based Deep Learning

**DOI:** 10.1101/2025.02.25.639900

**Authors:** Hoang-Hai Nguyen, Josip Rudar, Nathaniel Lesperance, Oksana Vernygora, Graham W. Taylor, Chad Laing, David Lapen, Carson K. Leung, Oliver Lung

**Affiliations:** National Centre for Foreign Animal Disease, Canadian Food Inspection Agency, Winnipeg, Manitoba, Canada; Department of Computer Science, University of Manitoba, Winnipeg, Manitoba, Canada; Department of Biological Sciences, University of Manitoba, Winnipeg, Manitoba, Canada; Department of Integrative Biology & Centre for Biodiversity Genomics, University of Guelph, Guelph, Ontario, Canada; School of Engineering, University of Guelph, Guelph, Ontario, Canada; Vector Institute, Toronto, Ontario, Canada; Agriculture and Agri-Food Canada, Ottawa, Canada

**Author notes:** Address correspondence to Hoang-Hai Nguyen and Josip Rudar.

**Keywords:** deep learning, alternate attention mechanisms, Fourier Transform, Wavelet Transform, chaos game representation, influenza A virus, antigenic types, viral host source, viral genomics

## Abstract

**Background:** Influenza A virus (IAV) poses a significant threat to animal health globally, with its ability to overcome species barriers and cause pandemics. Rapid and accurate IAV subtypes and host source prediction is crucial for effective surveillance and pandemic preparedness. Deep learning has emerged as a powerful tool for analyzing viral genomic sequences, offering new ways to uncover hidden patterns associated with viral characteristics and host adaptation.

**Findings:** We introduce WaveSeekerNet, a novel deep learning model for accurate and rapid prediction of IAV subtypes and host source. The model leverages attention-based mechanisms and efficient token mixing schemes, including the Fourier Transform and the Wavelet Transform, to capture intricate patterns within viral RNA and protein sequences. Extensive experiments on diverse datasets demonstrate WaveSeekerNet’s superior performance to existing models that use the traditional self-attention mechanism. Notably, WaveSeekerNet rivals VADR (Viral Annotation DefineR) in subtype prediction using the high-quality RNA sequences, achieving the maximum score of 1.0 on metrics including the Balanced Accuracy, F1-score (Macro Average), and Matthews Correlation Coefficient (MCC). Our approach to subtype and host source prediction also exceeds the pre-trained ESM-2 (Evolutionary Scale Modeling) models with respect to generalization performance and computational cost. Furthermore, WaveSeekerNet exhibits remarkable accuracy in distinguishing between human, avian, and other mammalian hosts. The ability of WaveSeekerNet to flag potential cross-species transmission events underscores its significant value for real-time surveillance and proactive pandemic preparedness efforts.

**Conclusions:** WaveSeekerNet’s superior performance, efficiency, and ability to flag potential cross-species transmission events highlight its potential for real-time surveillance and pandemic preparedness. This model represents a significant advancement in applying deep learning for IAV classification and holds promise for future epidemiological, veterinary studies, and public health interventions.

## 1 Introduction

Influenza viruses are a constant threat to public and animal health and a leading cause of acute respiratory diseases worldwide. Avian influenza is a contagious disease caused by infection with influenza A virus (IAV), which naturally resides and circulates in waterfowl [1]. Host barriers prevent IAV from freely infecting new non-avian hosts; however, cross-species transmission can occur when viruses evolve to overcome these barriers [2]. One crucial determinant of IAV infection is hemagglutinin (HA) receptor binding specificity. Human influenza virus strains prefer to bind to α2,6-sialic acid linkages, whereas avian virus strains preferentially bind receptors of α2,3-sialic acid linkages [3]. IAV has a complex genome structure with eight negative-sense, single-stranded RNA gene segments: PB2, PB1, PA, HA, NP, NA, M, and NS [4]. The hemagglutinin (HA) and neuraminidase (NA) gene segments are two major determinants of IAV antigenicity and virulence. IAV is classified into subtypes based on the HA and NA segments. There are 18 different HA subtypes (H1 through H18), 11 different NA subtypes (N1 through N11), and a possibility for new subtypes to be discovered, such as putative H19 [5], creating many possible combinations, such as H5N1, H3N2, and H7N9. Since 1918, there have been four major influenza epidemics and pandemics caused by IAV: the Spanish flu 1918 (H1N1), the Asian flu 1957 (H2N2), the Hong Kong flu 1968 (H3N2), and the 2009 swine flu pandemic (H1N1) [6,7]. Between 2003 and 2005, H5N1 IAV emerged in China and spread to other countries, causing widespread poultry outbreaks across Asia, Africa, the Middle East, and Europe [8,9]. In November 2021, a highly pathogenic avian influenza (HPAI) variant of H5N1 IAV emerged in domestic birds at a farm in Newfoundland, Canada, likely caused by transatlantic spread from Europe by wild birds [10]. Since its emergence, H5N1 IAV has circulated throughout North America. As of May 2025, this outbreak has affected more than 14.5 million birds in Canada [11] and more than 169 million birds in the United States [12]. This has significantly impacted poultry production, leading to rising egg prices and food inflation in the United States [13]. The virus has also spread to mammals such as skunks, foxes, raccoons, dogs, and dairy cows, including humans [14,15]. Because of this widespread impact and potential for further spread, understanding and predicting the host source of IAV is crucial for active surveillance and our preparedness for future pandemics. There is also a considerable need to accurately and rapidly identify different circulating IAV subtypes. Such identification is essential for effectively treating new and existing infections, as well as for planning and executing robust strategies for viral surveillance and outbreak prevention [16].

Although progress has been made in understanding IAV transmission, significant knowledge gaps remain [17,18]. Recently, machine learning (ML) and artificial intelligence (AI) have been used to analyze large-scale viral genomic sequences to facilitate a deeper understanding of the evolution and biology of pathogens. For example, machine learning algorithms like Random Forest and Support Vector Machine (SVM) have been used to identify genomic patterns associated with expanded host-range in severe acute respiratory syndrome coronavirus 2 (SARS-CoV-2) [19] and IAV [20–23]. However, increasing focus is being directed towards deep learning methods due to their robust applicability and ability to uncover complex relationships by weighing and transforming information contained within multiple genomic regions within a single model [24,25].

Ensemble learning has been successfully applied to supervised classification tasks in diverse fields [26,27]. Breiman (2001) demonstrated that this procedure is successful since decorrelation between ensemble members enables diverse learners to compensate for the errors of other ensemble members [28]. This observation was further developed in methods such as stacked generalization and voting classifiers, which exploit differences in the inductive biases of different machine learning algorithms to improve classification performance [29]. Deep learning has also leveraged the advantages of ensembles. For example, deep ensembles, which borrow from the work of Breiman (2001), have been used as an alternative to Bayesian Neural Networks to estimate prediction uncertainties [30,31]. Other innovations, such as dropout [32], act like ensembling by activating a different set of neurons with each training iteration. Finally, approaches such as multi-head attention [33] and Mixture-of-Experts [34] also contribute to good generalization.

This study presents a new deep learning model, WaveSeekerNet, based on an ensemble of different efficient attention-like and feed-forward network (FFN) mechanisms. WaveSeekerNet first splits image representations of RNA and protein sequences into different ‘word’ patches known as tokens. Our ensemble attention-like mechanism, the WaveSeeker block, then aggregates information from each token, forming a kind of genomic signature allowing the modelling of underlying biological patterns and functional relationships tied to important outcomes such as the host source. WaveSeekerNet precisely classifies subtypes and predicts the host source of IAV using the RNA and protein sequences of the HA and NA gene segments. Tests on held-out data reveal that WaveSeekerNet achieved state-of-the-art performance. Furthermore, WaveSeekerNet successfully identified zoonosis, reverse zoonosis, and enzootic strains.

## 2 Material and Methods

### 2.1 Influenza A virus dataset and general workflow

The RNA and protein sequences of the HA and NA segments were downloaded from EpiFlu GISAID in January and June 2024 [35] along with subtype, host, and other available metadata. The EpiFlu GISAID database contains numerous identical sequences (Figure 1a). For example, the HA segment of the avian H5N1 strain *A/goose/Zhejiang/727098/2014* (EPI681274) is identical to that of the swine H5N1 strain *A/swine/Zhejiang/SW57/2015* (EPI1600724). Effective deduplication, particularly across different hosts, presents challenges. In this work, we kept the earliest collected sequence and its associated metadata, based on the inference that this sequence originated from the initially identified host. Any identical sequences that were collected at a later time were removed from the analysis. After this step, an additional length and ambiguity filtering was performed (Figure 1a). The lengths of the HA and NA segments from the reference strain *A/New York/392/2004* (H3N2 - HA segment: EPI79008, NA segment: EPI79013) were used as the baselines for identifying sequences that are either too short or too long. HA and NA sequences with a length at least 80% and at most 120% of the reference and with a maximum of 10 ambiguous sequence characters were included in the set of high-quality sequences. Sequences that did not meet either of these criteria were placed into a low-quality dataset, and if they had an extended minimum length of 1000nt or 350aa (for HA) and 850nt or 250aa (for NA). Otherwise, they were excluded from the analysis (Figure 1a). We screened RNA sequences in the collected datasets using the VADR (Viral Annotation DefineR) tool [36] and reverse-complemented any sequences not in the 5’ to 3’ direction.

**Figure 1:**
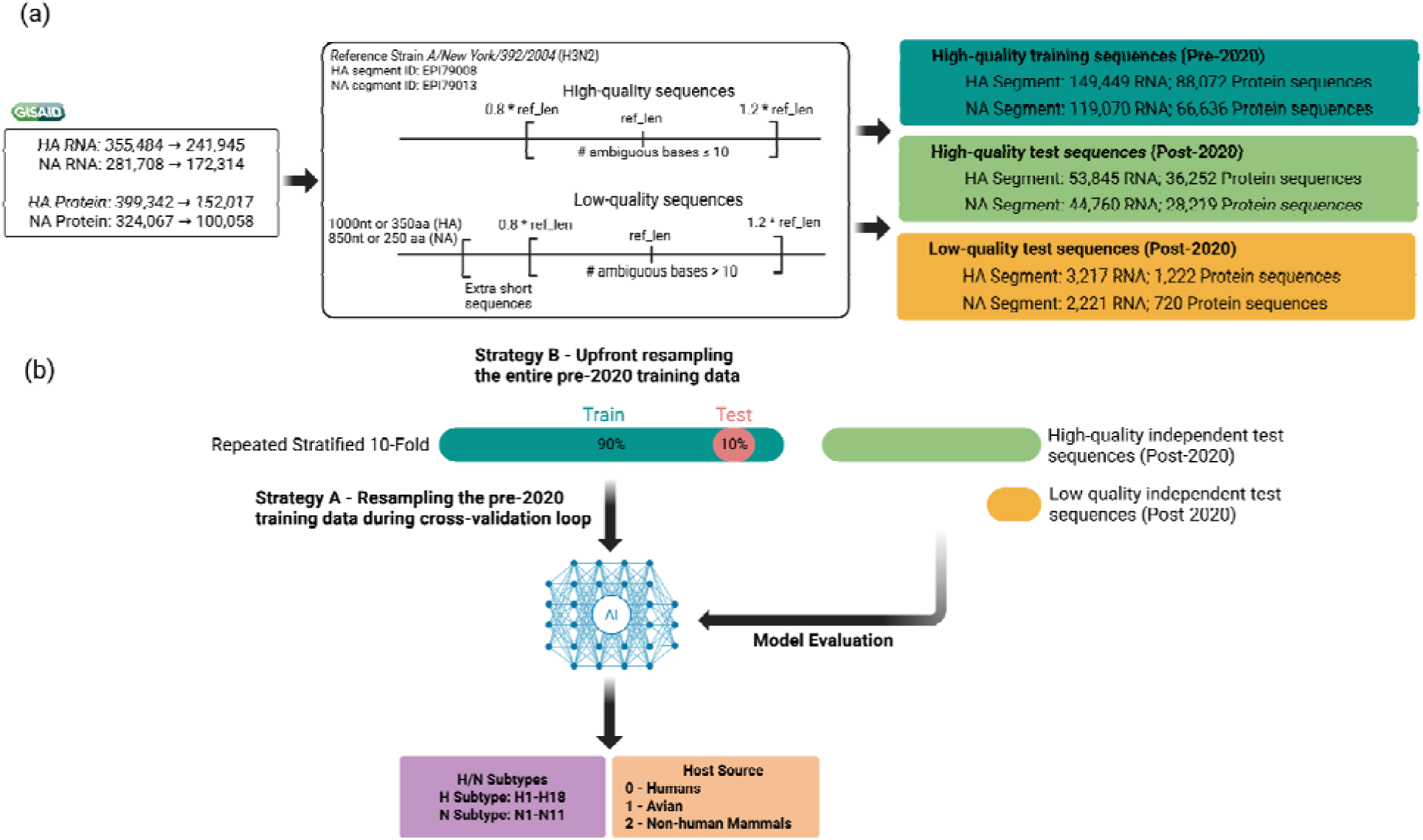
The general workflow consists of several steps. (a) The IAV sequences used in this study were retrieved from EpiFlu GISAID. The subtype information of each sequence and the host source information from which each sequence was obtained were also downloaded. If two or more sequences were identical, only one sequence with the earliest collection date and associated metadata was retained. Next, the sequences underwent quality control and binning into training and test sets. (b) The quality-controlled sequences were encoded into the form of images (2D matrix) using one-hot encoding and Frequency Chaos Game Representation (FCGR). Repeated stratified 10-fold cross-validation was used to train each model. Two training data resampling strategies were deployed to evaluate the models: (Strategy A) resampling the pre-2020 training data within each cross-validation fold versus (Strategy B) resampling the entire pre-2020 training data before cross-validation. High-quality and low-quality independent test sets were used to assess the overall quality of the trained models. This figure was created with BioRender.

To predict future viral characteristics, we temporally split the high-quality and low-quality sequences. High-quality sequences collected before January 1, 2020, were reserved for training, while those collected since 2020 were used for evaluating generalization performance (Figure 1a). This strategy ensured that highly similar strains were not found in training and test sets. Due to the over-representation of some HA and NA subtypes and the under-representation of others, we down-sampled over-represented subtypes (defined as having more than 6,000 sequences) to 6,000 sequences per subtype. Examples of over-represented subtypes include H1 (56,593 RNA sequences), H3 (59,661 RNA sequences), N1 (47,592 RNA sequences), and N2 (58,750 RNA sequences). Rare subtypes, defined as having fewer than 600 sequences, were up-sampled to 600 sequences per subtype. Notably, subtypes H17 (2 sequences), H18 (1 sequence), N10 (2 sequences), and N11 (1 sequence) are extremely rare and were only included in the training data (Supplementary Tables S1, S3). Maintaining sequence diversity during training is crucial to prevent model bias, particularly towards rare subtypes. While the approximately 6,000-sequence provides sufficient diversity for prevalent subtypes [37], the 600-sequence for rare subtypes requires up-sampling to mitigate this risk. We performed down-sampling and up-sampling (copying the original multiple times) using the ‘resample’ function from scikit-learn (v.1.5.1). This resulted in a final training set consisting of 18 HA and 11 NA subtypes (classes) (Figure 1b).

Sequences were also labeled for the host source prediction datasets, according to the source from which they were isolated. In this work, we grouped sequences into three major host groups: Humans, Avian (e.g., falcon, turkey, goose), and Non-human Mammals (e.g., swine, horses, dogs) (Figure 1b). Since the human and avian hosts accounted for the majority of sequences, these groups were down-sampled to 16,000 sequences per group, while all Non-human mammal sequences were used. We used the high-quality and low-quality sequence collected after December 31, 2019, to evaluate the performance of the models. We also identified the subset of strains where both an HA and NA sequence were present and used these sequences together for host source prediction (Figure 2a and Supplementary Table S5). The data distribution used in this study is shown in Supplementary Tables S1-5, and Figure 1 illustrates the major steps in this study.

**Figure 2:**
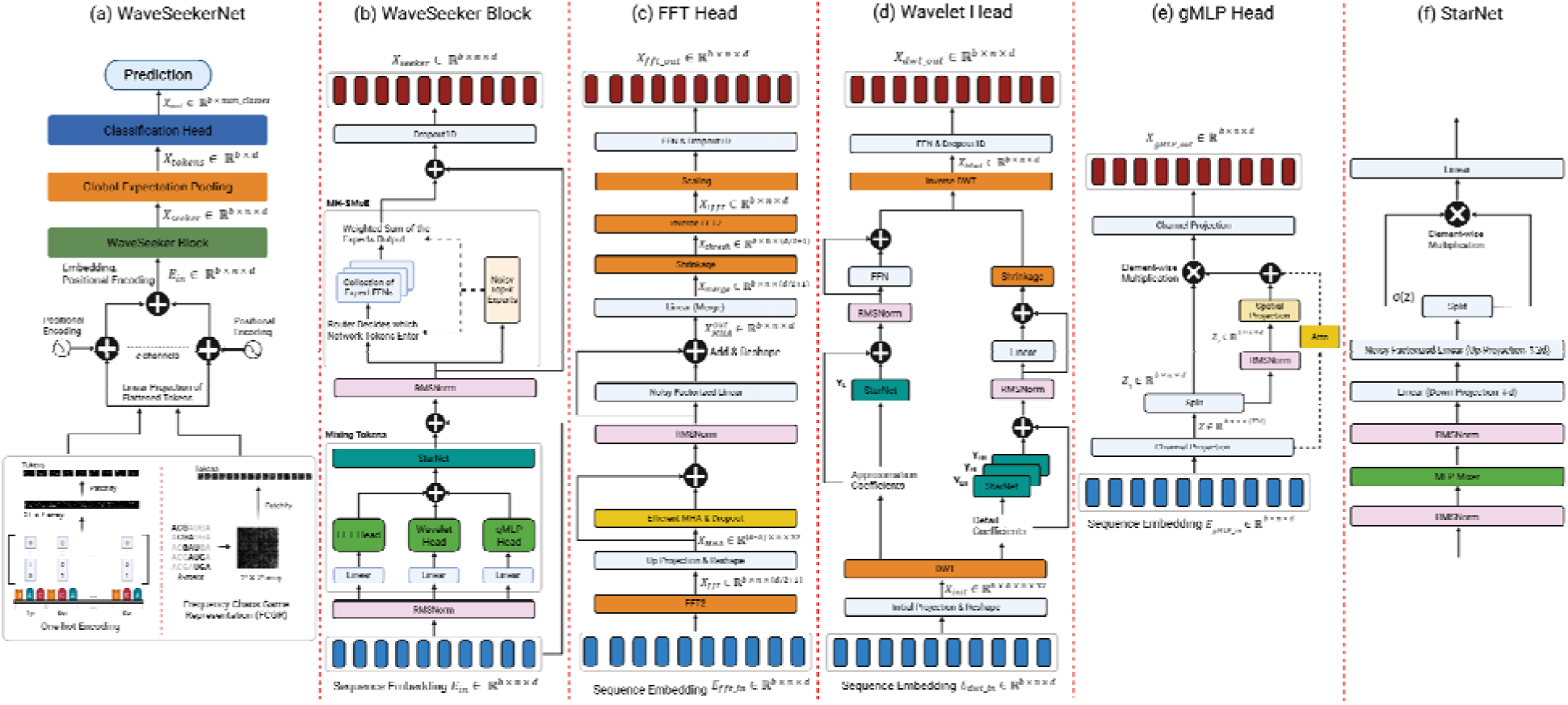
(a) The overall structure of WaveSeekerNet. (b) The WaveSeeker block contains token mixing schemes based on (c) the Fourier Transform, (d) the Wavelet Transform, and (e) gMLP. A modified StarNet (f) that uses an MLP-Mixer layer and noisy factorized linear layers is employed following token mixing. This is followed by an MH-SMoE layer, which processes tokens along the hidden dimension. This figure was created with BioRender.

### 2.2 Transforming sequences into feature vectors

To enable the neural network models to learn and recognize patterns of RNA and protein sequences, we employed two methods to encode sequences into the form of images: Frequency Chaos Game Representation (FCGR) and one-hot encoding. The information content within nucleic acid and protein sequences can be viewed as a signal that can be transformed into an image and used as input for machine learning algorithms [38–40]. Numerous examples in the literature have taken the Chaos Game Representation (CGR) approach. In the early 1990s, for example, the CGR of nucleic acids was used to visualize the structure of DNA sequences [41]. The CGR maps a DNA/RNA sequence *s* onto the unit square using an algorithm illustrated in Supplementary Algorithm S1.

An extension of CGR, the FCGR [41–43], subdivides the CGR into a 2^*k*^ × 2^*k*^ grid of sub-squares. Each sub-square corresponds uniquely to a particular *k*-mer and represents its number of occurrences (i.e., its frequency) within the original sequence *S*. Due to variations in sequence length within the collected data, *k*-mer frequencies within the FCGRs require standardization for comparison. To address this, we applied the standardization method proposed by Wang et al. [44] (Equation 1) to create the standardized FCGR, *Ā*, which was used for our analysis of IAV RNA sequences. FCGRs were created using the ‘complexCGR’ package version 0.8.0 in a Python 3.12 environment [45].

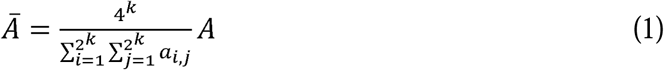

where *A* is the *k*th-order FCGR of the sequence *S*, and *a_i,j_* denotes the elements of *A*.

We used one-hot encoding for protein sequences to transform sequences into representations suitable for machine learning applications. This method has been used successfully to study influenza viruses, rotavirus A, and rabies lyssavirus and to predict host-range, antigenic types, and pathogenicity [25,46]. Each character in the protein sequence is encoded as a 21-dimensional vector via one-hot encoding. The aggregation of these vectors forms a 2D matrix encoding an amino-acid sequence. For example, A (Alanine) can be encoded as (1, 0, 0, …, 0, 0, 0), C (Cysteine) as (0, 1, 0, …, 0, 0, 0), D (Aspartate) as (0, 0, 1, …, 0, 0, 0), and X as (0, 0, 0, …, 0, 0, 1). We zero-padded the C-terminal end of each HA and NA polypeptide sequence to ensure that the inputs to each deep learning model have the same length.

## 3 Model Architecture

We propose a deep learning network called WaveSeekerNet, as illustrated in Figure 2, which is based on the attention-like architecture [33] and Vision Transformer [47]. Figure 2a presents the overall structure of WaveSeekerNet. The first step of our approach is to split the FCGRs and one-hot encodings of input sequences into ‘word’ patches (tokens). These ‘word’ patches are then flattened and linearly projected using noisy factorized linear layers to create an embedding of each patch *E_patch_* [48]. A learnable sinusoidal positional encoding, based on positional encoding T in the Transformer model, is then added to Ii_rpatch_ according to Equation 2 to create the final embedding *E_in_* ∈ ℝ^*b* × *n* × *d*^ [49], where *b*, *n*, and *d* are the batch, token, and hidden (embedding) size, respectively. For multiple channels, WaveSeekerNet concatenates the embeddings along the token dimension. The final embedding *E_in_* is fed to the transformer-like WaveSeeker block (Figure 2b). After passing through the WaveSeeker block, the transformed embeddings are pooled using Global Expectation Pooling [50] and sent to the classification head. Finally, the traditional feed-forward network or the Kolmogorov-Arnold Network (KAN) [51] will perform the classification.

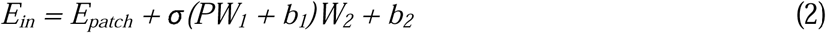

### 3.1 WaveSeeker block

The key component of WaveSeekerNet is the WaveSeeker block (Figure 2b). We incorporate significant changes in this block to utilize lessons learned from ensemble approaches. In this study, the attention mechanism [33] is replaced by three token mixing approaches: the Fourier Transform (Figure 2c), the Wavelet Transform (Figure 2d), and th gating Multilayer Perceptron (gMLP) architecture (Figure 2e) [52]. The outputs of these token mixing blocks are concatenated along the hidden dimension and then merged using a modified version of the recently released StarNet [53]. In the StarNet block (Figure 2f), we replace depthwise convolution with an MLP-Mixer layer [54] and apply noisy factorized linear layer [48] before the star operation (element-wise multiplication). To improve the network capacity, the traditional feed-forward layer of Vision Transformer [47] is replaced by a Sparsely Gated Multi-Head Mixture-of-Experts layer (MH-SMoE) [34,55,56].

#### 3.1.1 Fourier Transform block

A significant amount of effort has been directed towards developing alternatives to the self-attention mechanism due to its high computational and memory cost [54,57,58]. Research conducted thus far strongly suggests that algorithms that efficiently share information between tokens are required to develop an efficient alternative to the transformer; one of the first works to identify this approach is the FNET [57]. At the heart of this alternative to the transformer is the Fourier Transform. FNET applies a 2D discrete Fourier Transform (DFT) to the input embeddings. This transform block mixes information from each of the tokens, and during training, the model learns the weights associated with the best combination of tokens that minimizes the loss function. In image processing, the Fast Fourier Transform (FFT) is commonly used to compute the DFT, transforming an image into its frequency domain. Low frequencies represent global patterns, while high frequencies correspond to abrupt changes, such as edges, providing more image details. The 2D DFT and inverse DFT (iDFT) of an image array can be calculated using Equations 3 and 4:

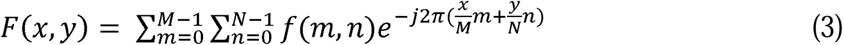

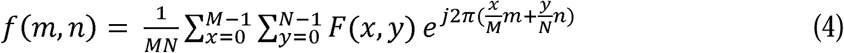

where *F*(*x*, *y*) is the function representing the image in the frequency domain, *f*(*m*, *n*) is a pixel at position (*m*, *n*) in the spatial domain, and *M* × *N* represents the image’s dimensions.

In this study, we modified the original FNET architecture [57] to design an additional token mixing scheme as illustrated in Figure 2c. Tokens are first mixed by applying the 2D DFT to transform the sequence embedding 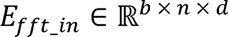 into frequency coefficients. Like FNET, we only keep the real part of the transform result. Since the FFT of a real signal is Hermitian-symmetric, we omit the negative frequencies, producing 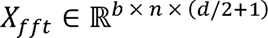. Subsequently, *X*_*fft*_ is projected back to the embedding dimension *d* and reshaped to create multi-head 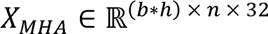, where the number of heads *h* = *d*/32. We then employ an efficient attention with linear complexities [58] on *X_MHA_* to capture intricate interactions within the frequency space. The output of this efficient attention mechanism is then enhanced with dropout, RMS normalization [59], and skip connections, yielding 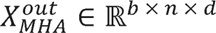. Next, we merge heads of 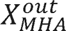 into 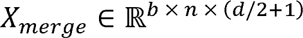. This merging prepares the representation for subsequent processing and transformation back into the spatial domain.

Likely, the information needed to reconstruct each embedding from *X_merge_* using the iDFT will be concentrated within the low-frequency components, presenting an opportunity to promote sparsity and regularize this block of the network. This can be accomplished by applying a soft-thresholding operation on *X_merge_* (Equations 5-7) [60]. Finally, tokens are demixed by inverse FFT (iFFT), which transforms the thresholded *X_thresh_* (Equation 7) back into the spatial domain 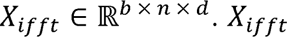 is then scaled so that each element is between +5 and -5, which is then passed through the feed-forward network, followed by dropout along the token dimension. Supplementary Algorithm S2 presents the pseudo-code of the Fourier Transform block.

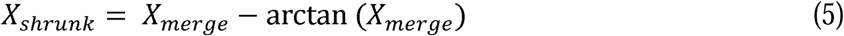

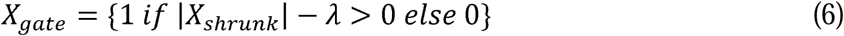

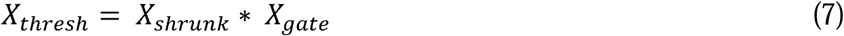

#### 3.1.2 Wavelet Transform block

While the Fourier Transform captures global frequency information, it does not localize the frequency within a sequence. The Wavelet Transform can address this shortcoming since this transformation can identify where frequency components occur within a signal. This capacity is particularly useful in sequence analysis since biological signals found in nucleic acid and protein sequences change over time [61]; thus, the Wavelet Transform can be applied to locate relevant frequency components in local regions of the sequence. In image processing, the discrete Wavelet Transform (DWT) is used to divide spatial information present in the image into low-frequency and high-frequency components corresponding to approximation and detail coefficients, respectively. In the Wavelet Transform, a signal is convolved with bandpass filters or mother wavelets *ψ_a,b_*(*t*) (Equation 8), where *a* and *b* determine the scale and location of the wavelet, respectively [62,63]. The wavelet will be compressed when the scaling value *a* decreases, capturing high-frequency components. In contrast, increasing the scaling value *a* will stretch the wavelet and capture low-frequency components. The translational value *b* shifts the wavelet in time (or space), allowing us to localize where specific frequency components occur in the signal.

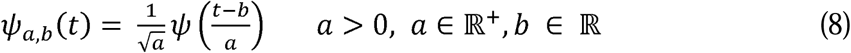

Formally, let 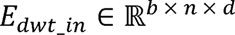 be the sequence embedding, we perform initial projection on *E_dwt_in_* and then split the embedding dimension into *h* heads, where *h* = *d*/32, creating initial input 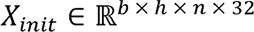 for the Wavelet Transform. Next, we apply 2D DWT using the Pytorch Wavelets package version 1.3.0 [64] to transform the input space *X_init_* into the Wavelet coefficients. Concretely, the Wavelet Transform applies low-pass and high-pass filters to transform *X_init_* into *Y_L_* and *Y_H_* subbands. *Y_L_* refers to the approximation coefficients that reflect the overall structure of the input space at a coarse-grained level. *Y_H_* represents detail coefficients at a fine-grained level, which is a single stacked tensor of *Y_LH_* (horizontal detail), *Y_HL_* (vertical detail), and *Y_HH_* (diagonal detail). Rather than employing an efficient attention layer (which was utilized in the Fourier Transform block), the Wavelet coefficients *Y_L_* and *Y_H_* are processed by StarNet layers. Like the Fourier Transform block, we apply the skip-connections and RMS normalization in the Wavelet coefficient space and shrink high-frequency components in the *Y_H_* subbands using Equations 5-7. The processed *Y_L_* and *Y_H_* are then used to transform feature maps back into the spatial domain using inverse DWT (iDWT), which is then passed through a feed-forward network, followed by dropout along the token dimension. Supplementary Algorithm S3 presents the pseudo-code of the Wavelet Transform block.

## 4 Implementation and evaluation methods

### 4.1 Model selection, training, and testing

In this study, we compared the prediction performance of WaveSeekerNet with that of established Transformer-only models using the traditional self-attention mechanism. Xu et al. previously employed the Transformer-only models to predict IAV host source and antigenic types [24,65]. These models were among the most effective applications of machine learning algorithms for predicting IAV host source and subtypes. As part of our comparative analysis, we explored the integration of the FNET architecture [57] into the Transformer-only models, replacing the conventional multi-head self-attention mechanism. We also evaluated the performance of the pre-trained ESM-2 models with transfer learning for protein sequences [66]. Table 1 presents the hyperparameter settings used to train deep learning models. We also introduce a new activation function, ErMish (Equation 9), which allows for a greater range of negative activations. This activation function uses a learnable parameter, *α*, to adjust how positive and negative input values influence the magnitude and sign of the function’s output:

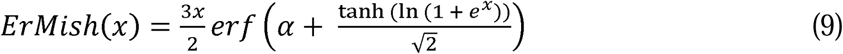

**Table 1:** The hyperparameter settings used by the WaveSeekerNet, Transformer-only, and pre-trained ESM-2 models during cross-validation.

| Model | Hyperparameter Settings | Explanation of Parameters |
| --- | --- | --- |
| WaveSeekerNet | Default Settings (Baseline Model):<br>*use_fft* = True<br>*use_wavelets* = True<br>*use_gmlp* = True<br>*wavelet_names* = “sym4”<br>*emb_dim* = 64<br>*final_hidden_size* = 24<br>*use_kan* = True<br>*use_smo* = True<br>*use_gc* = True<br>*use_lookahead* = True<br>*activation* = ErMish<br>Common Settings:<br>*batch_size* = 256<br>*epochs* = 35<br>Settings for FCGR representation of RNA sequences (apply for all hyperparameters, models):<br>*k-mer* = 6 (FCGR array size of $64 \times 64$)<br>*patch_size* = (4, 4)<br>Settings for one-hot encoding data of protein sequences (apply for all hyperparameters, models):<br>*patch_size* = (3, 21) | *use_fft* – Specify whether the Fourier Transform block is used in the WaveSeeker block.<br>*use_wavelets* – Specify whether the Wavelet Transform block is used in the WaveSeeker block.<br>*use_gmlp* – Specify whether the gMLP block is used in the WaveSeeker block.<br>*wavelet_names* – Specify the wavelet family for the Wavelet Transform. It will be used when *use_wavelets* = True.<br>*emb_dim* – The embedding dimension of the model.<br>*final_hidden_size* – The size of the penultimate layer in the classification head.<br>*use_kan* – Specify whether the KAN network is used as the classification head.<br>*use_smo* – Specify whether the MH-SMoE network is used in the FFN layer of the WaveSeeker block.<br>*use_gc* – Specify whether the Gradient Centralization is used in the optimizer [70].<br>*use_lookahead* – Specify whether lookahead is used in the optimizer [71].<br>*activation* – Specify activation for model (ErMish, Mish [72], GELU [73], ReLU [74]).<br>*k-mer* – the *k*th-order of FCGR.<br>*patch_size* – size of ‘word’ patches. |
| Pre-trained ESM-2 (esm2_t6_8M_UR50D or we called ESM-2_t6 for short) | *n_layers* = 6<br>*emb_dim* = 320<br>*batch_size* = 64 | *n_layers* – the number of the Transformer encoder layers.<br>*emb_dim* – The embedding dimension of the model. |
| Pre-trained ESM-2<br>(esm2_t12_35M_UR50D<br>or we called ESM-2_t12<br>for short) | • *n_layers* = 12<br>• *emb_dim* = 480<br>• *batch size* = 32 | • *n_layers* – the number of the Transformer encoder layers.<br>• *emb_dim* – The embedding dimension of the model. |
| Transformer-only using<br>FNET | • *emb_dim* = 64, 128<br>• *activation* = ReLU<br>• *epochs* = 150<br>• *batch size* = 256 | • *emb_dim* – The embedding dimension of the model. |
| Transformer-only using<br>conventional multi-head<br>attention (MHA) | • *emb_dim* = 64, 128<br>• *nhead* = 4<br>• *activation* = ReLU<br>• *epochs* = 150<br>• *batch size* = 256 | • *emb_dim* – The embedding dimension of the model.<br>• *nhead* – The number of heads in the multi-head attention. |

We trained WaveSeekerNet using a composite loss function (Equation 10). The first part of the composite loss function is the cross-entropy loss function. This function is typically used to measure how much the prediction from the model, *ŷ*, deviates from the expected classification outcome, *y*. The second part of the loss function, the router z-loss, penalizes large logits during routing into the MoE network to force the model to balance the number of tokens routed to each expert. The last part of the loss function encourages the selection of specific activation functions within each of the KAN networks used in the classification head by reducing the impact of unnecessary activation functions on the final output of the KAN layer [51]. See Supplementary Algorithms S4 and S5 for details.

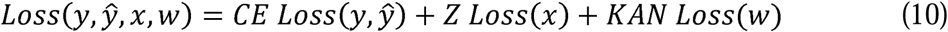

We observed that the Transformer-only models required more converging time than our model. Therefore, the Transformer-only models were trained with 150 epochs, while WaveSeekerNet and the pre-trained ESM-2 models were trained with only 35 epochs. We used repeated stratified 10-fold cross-validation to assess the generalization performance of models (Figure 1b). The test set within each cross-validation fold was used to check and ensure the training process functioned properly. The early-stopping-pytorch package [67], with a patience value set to 7 and a delta of 0.001, was used to determine when the pre-trained ESM-2 models converged, and training could cease. All models were trained on an NVIDIA GPU A100. For model evaluation, the weights of the last epoch (except for the pre-trained ESM-2 models that use the early stopping strategy) were used. The reported scores represent the mean across the 10 folds. Finally, a negative control is needed for a proper assessment of learning. The dummy classifier from scikit-learn (v.1.5.1) using a stratified selection strategy was used for this purpose.

We conducted ablation studies to evaluate the impact of different hyperparameters (Table 1) on WaveSeekerNet’s performance. We used the baycomp package version 1.0.3 [68,69] to compare the generalization performance of WaveSeekerNet with various hyperparameters. In this work, we performed up-sampling for the minority classes and down-sampling for the majority classes in the pre-2020 training data during cross-validation (Strategy A). In this resampling approach, we performed up-sampling for the sporadic subtypes, which have fewer than 10 sequences, such as H15 (5 protein sequences), H17 (2 sequences), H18 (1 sequence), N10 (2 sequences), and N11 (1 sequence) before cross-validation. This procedure is necessary to maintain the diversity and the same number of classes for training data in each cross-validation fold; otherwise, the sporadic subtypes would only be seen in a single or several folds. We also tested the second scenario of resampling, in which up-sampling and down-sampling were performed on the entire pre-2020 dataset before cross-validation (Strategy B). This was done to evaluate how up-sampling and down-sampling impact bias in predicting host source and subtypes. The ‘two_on_single’ function in the baycomp package was used to determine the probability of one scenario being better. To do this, we chose a region of practical equivalence (ROPE) to be 0.025. This region quantifies the probability that the overall generalization performance of models of each resampling approach differs by less than 0.025.

### 4.2 Sequence similarity search methods

We evaluated the performance of deep learning models against BLAST, a widely used sequence similarity search method in computational biology and bioinformatics. Comparisons to the BLAST search served as a sanity check to ensure that our model and other deep learning methods did not suffer from systemic failures and worked as expected. BLASTp (v2.16.0) [75] with options ‘-num_alignments 100 -evalue 1e-5’ was used for protein sequence-based subtype prediction. The BLASTp database was built using the same training splits of the pre-2020 high-quality dataset used for model training. This was done to ensure that results using sequence similarity search methods would be comparable. The query is the sequences in the post-2020 high-quality and low-quality test data. The top hit was determined to be the hit with the highest bit score after filtering for hits with at least 85% identity and an alignment length of at least 50. We assigned an arbitrary subtype (1-18 for HA subtype, 1-11 for NA subtype) as the prediction to query sequences without hits.

We used VADR [36], a viral sequence classification and annotation tool for subtype prediction using RNA sequences. VADR uses a model library from a subset of references to classify and annotate virus sequences. We used the *v-annotate.pl* VADR script (v1.6.4) with influenza models (v1.6.3-2) [76] for subtype classification.

### 4.3 Analysis of the ongoing H5Nx avian influenza outbreak in North America

To investigate WaveSeekerNet’s ability to identify the most probable source of a transmission event, we used data from the ongoing H5Nx avian influenza outbreak in North America as a test case. We identified a subset of our data consisting of 1,659 unique HA RNA sequences from various H5Nx strains circulating in North America in 2023 and 2024 [35]. These strains represent a recent snapshot of the ongoing H5Nx avian influenza outbreak. To support the predictions made by WaveSeekerNet, we performed a phylogenetic analysis of the H5Nx virus dataset. Sequences were aligned with MAFFT (RRID:SCR_011811, v7.520) [77,78]. Time-calibrated phylogenetic analysis was performed in BEAST (RRID:SCR_010228, v1.10.4) [79] using a coalescent Bayesian skyline model [80,81] and a general time-reversible substitution model with the across-site rate heterogeneity sampled from a gamma distribution with four discrete categories and an uncorrelated relaxed log-normal molecular clock [82]. Two independent runs were performed with a chain length of 50 million generations each, and parameter values were sampled every 1,000 generations. Stationarity and convergence of independent runs were assessed in Tracer v1.7 [83]. The LogCombiner module in the BEAST software was used to remove the burn-in fraction and combine the log and tree files. The maximum clade credibility tree with median node heights was produced using TreeAnnotator [79]. Phylogenetic trees were visualized using FigTree (RRID:SCR_008515, v.1.4.3) [84] and Inkscape [85].

### 4.4 Evaluation Metrics

Evaluation metrics used in this study include F1-score (Macro Average), Balanced Accuracy, and Matthews Correlation Coefficient (MCC). Since the dataset is imbalanced and we treat all classes equally regardless of their support values, we used the F1-score (Macro Average) to compare the performance between methods. This metric is computed by taking the arithmetic mean of the F1-score (Equation 11) of all classes. This measurement approach is appropriate since we are interested in assessing the performance across the diverse range of IAV subtypes and host categories in our data.

The equations of F1-score for each class, Balanced Accuracy, and MCC are defined as follows:

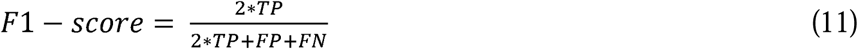

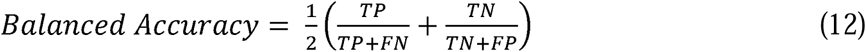

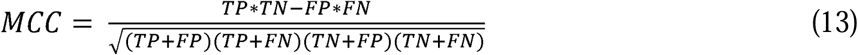

where *TP, FP, TN* and *FN* stand for True Positive, False Positive, True Negative and False Negative, respectively.

## 5 Results

### 5.1 WaveSeekerNet is robust to changes in hyperparameter settings

We conducted initial ablation studies to evaluate the impact of various hyperparameters, as illustrated in Table 1, on WaveSeekerNet’s performance. For example, we disabled one or two token mixing schemes and used various activation functions. The experimental results of host source prediction, presented in Supplementary Tables S6, S7 and Supplementary Figures S1, S2, S3, demonstrate WaveSeekerNet’s robustness to changes in hyperparameter settings under this testing regime. While some hyperparameter choices did influence performance, the baseline WaveSeekerNet model consistently demonstrated strong performance. This model of choice, using all three token mixing schemes (Fourier Transform, Wavelet Transform and gMLP blocks), ranked at or near the top across various data representations and datasets. When the baseline model did rank lower, the generalization performance of the alternative models was often similar to that of the baseline. Our results also show that disabling any of the three token mixing schemes decreased generalization performance in most RNA sequence tests, while yielding mixed results in protein sequence tests. Generally, empirical evidence from these tests indicated that the probability of modifications to the baseline settings is unlikely to improve classification performance (Supplementary Tables S6, S7).

WaveSeekerNet’s performance with the Wavelet Transform block was particularly impactful when trained on FCGR representation of RNA sequences (Supplementary Table S6 and Supplementary Figures S1a, S1b, S2a, S2b, S3a, S3b). The Wavelet Transform block improved the WaveSeekerNet’s performance significantly. For example, when tested with HA RNA sequences, using gMLP-only, FFT and gMLP, and FFT-only token mixing schemes, WaveSeekerNet achieved F1-scores (Macro Average) of 0.737±0.07, 0.824±0.084, and 0.781±0.121, respectively (Supplementary Table S6). While the baseline model (using all three token mixing schemes), Wavelet Transform-only, Wavelet Transform and gMLP, and FFT and Wavelet Transform achieved F1-scores (Macro Average) of 0.965±0.015, 0.962±0.033, 0.964±0.014, and 0.969±0.007, respectively.

The Lookahead optimizer was another hyperparameter that influenced generalization performance. Disabling this optimizer consistently degraded WaveSeekerNet’s performance across various data representations and datasets (Supplementary Tables S6, S7 and Supplementary Figures S1, S2, S3). For instance, when assessing the host source prediction using the FCGRs of the HA segment (Supplementary Table S6), the F1-scores (Macro Average) were 0.867±0.266 and 0.701±0.23 for high-quality and low-quality datasets, respectively, when the Lookahead optimizer was disabled. In the baseline model, where the Lookahead optimizer is enabled, the F1-scores (Macro Average) in these tests were 0.965±0.015 and 0.713±0.074, respectively. Gradient centralization was also important, but primarily impacted results in tests using FCGRs. Disabling gradient centralization resulted in up to an 11% decrease in the F1-score (Macro Average) (Supplementary Table S6 and Supplementary Figure S1a). Our use of the ErMish activation positively impacted WaveSeekerNet’s performance (Supplementary Tables S6, S7). Alternative activations, such as Mish, GELU, and ReLU, often had a probability of less than 0.5 of outperforming the baseline model. Finally, we observed that alternative configurations, such as disabling KAN and MH-SMoE, negatively impacted generalization performance. This observation was evident in both RNA and protein sequence datasets, where the baseline model without KAN or MH-SMoE generally underperformed the baseline model (Supplementary Tables S6, S7).

### 5.2 WaveSeekerNet achieves excellent subtype and host source classification using a parameter-efficient design

WaveSeekerNet also demonstrated a favorable balance between predictive performance and computational resource requirements. Due to the very high GPU memory requirements of the pre-trained ESM-2 model, we were restricted to a training batch size of 64 (for the ESM-2_t6 model) or 32 (for the ESM-2_t12 model). Furthermore, this approach demanded significantly more training time and parameters than WaveSeekerNet and Transformer-only, yet it only resulted in comparable generalization performance to WaveSeekerNet when tested on the high-quality protein sequences (Table 2 and Supplementary Table S8). When tested on the low-quality protein sequences, the performance of the pre-trained ESM-2 models was better than that of the Transformer-only models. However, the F1-score (Macro Average) of the pre-trained ESM-2 models was up to 46 percentage points lower than WaveSeekerNet’s score in these tests. For example, with HA subtype prediction using the low-quality protein sequences, the ESM-2_t12 model achieved an F1-score (Macro Average) of 0.473, while WaveSeekerNet with FFT-based head achieved an F1-score (Macro Average) of 0.932 (Table 2). Similarly, WaveSeekerNet with FFT-based and the ESM-2_t6 model achieved F1-scores (Macro Average) of 0.823 and 0.684, respectively, for host source prediction using the low-quality protein sequences of the HA segment (Supplementary Table S8). The FNET and MHA approaches took the least training time per epoch, but neither method could extract enough information to match the generalization performance of WaveSeekerNet. Finally, depending on the selected hyperparameters, WaveSeekerNet’s parameter count, and by proxy memory efficiency, can be on the same order of magnitude as the MHA-based transformer. For example, by only using the FFT-based head or disabling the KAN network in the classification head, total parameters are reduced by nearly 29% and 50%, respectively (Table 2). Notably, such changes do not meaningfully change the classification performance on the high-quality protein sequences.

**Table 2:**
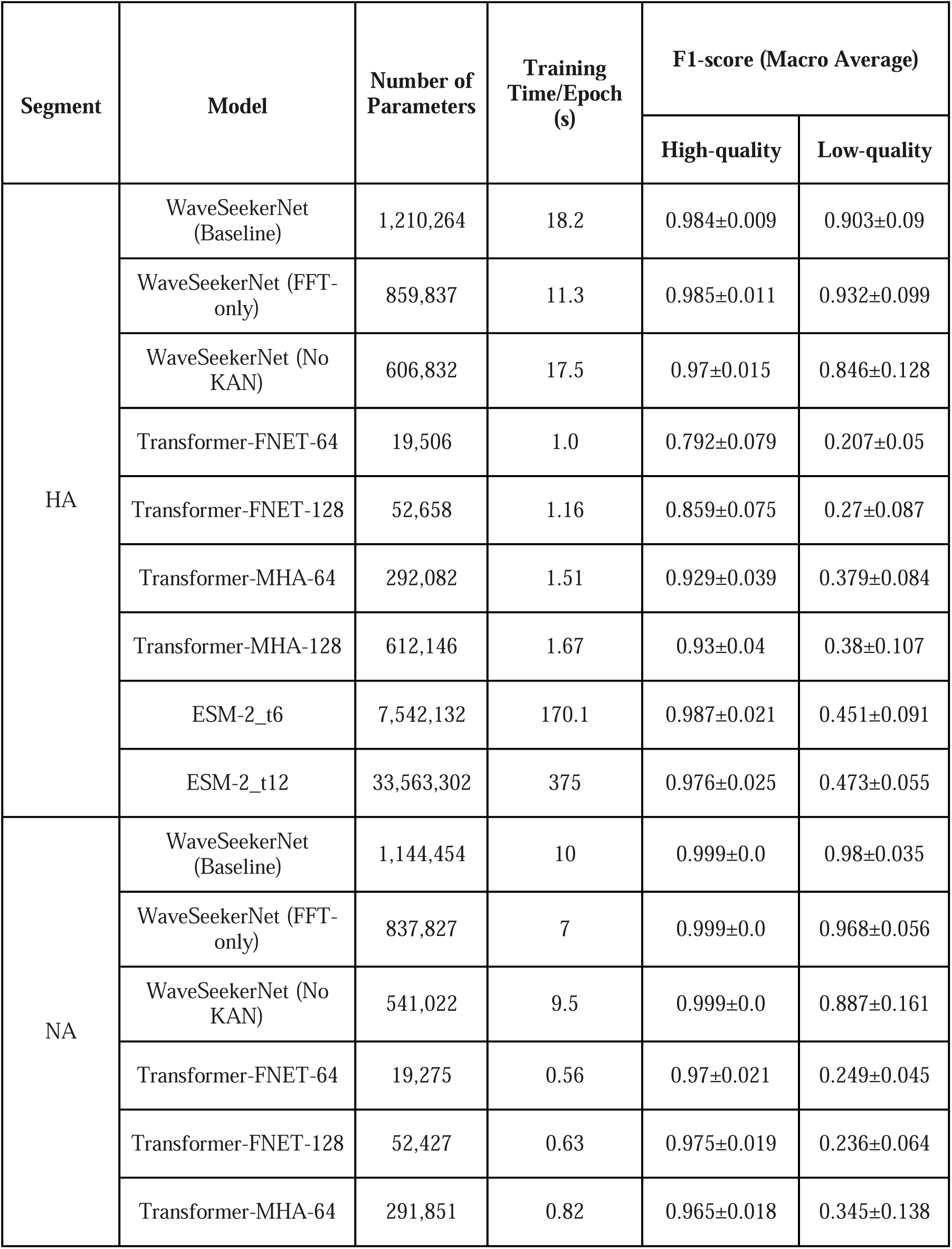
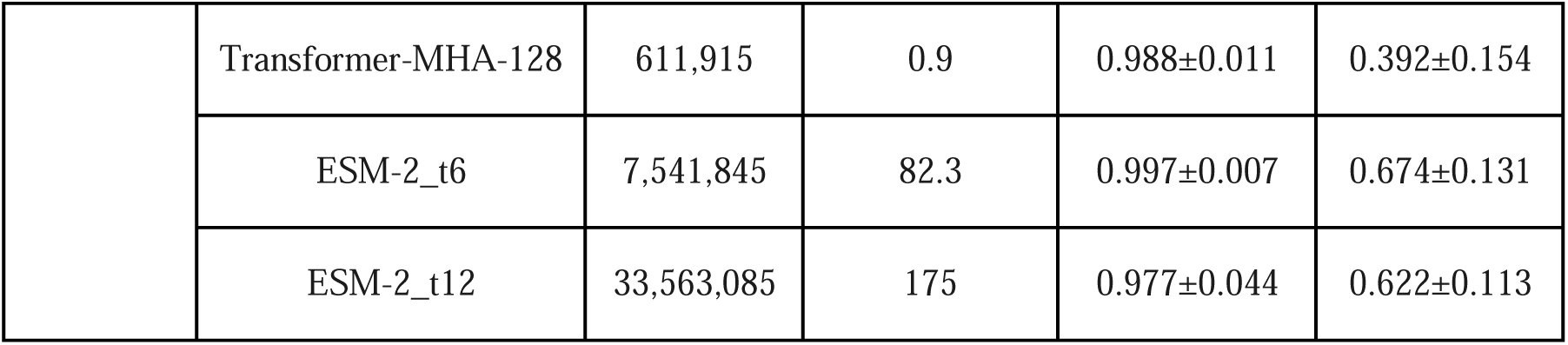
Comparison of computational costs and efficiency for the WaveSeekerNet, Transformer-only, and pre-trained ESM-2 models when trained to predict HA and NA subtypes using protein sequences. For the WaveSeekerNet models, we show the baseline model with complete options and the modified baseline models using the fewest parameters.

| Segment | Model | Number of Parameters | Training Time/Epoch (s) | F1-score (Macro Average) |  |
| --- | --- | --- | --- | --- | --- |
|  |  |  |  | High-quality | Low-quality |
| HA | WaveSeekerNet (Baseline) | 1,210,264 | 18.2 | 0.984±0.009 | 0.903±0.09 |
|  | WaveSeekerNet (FFT-only) | 859,837 | 11.3 | 0.985±0.011 | 0.932±0.099 |
|  | WaveSeekerNet (No KAN) | 606,832 | 17.5 | 0.97±0.015 | 0.846±0.128 |
|  | Transformer-FNET-64 | 19,506 | 1.0 | 0.792±0.079 | 0.207±0.05 |
|  | Transformer-FNET-128 | 52,658 | 1.16 | 0.859±0.075 | 0.27±0.087 |
|  | Transformer-MHA-64 | 292,082 | 1.51 | 0.929±0.039 | 0.379±0.084 |
|  | Transformer-MHA-128 | 612,146 | 1.67 | 0.93±0.04 | 0.38±0.107 |
|  | ESM-2_t6 | 7,542,132 | 170.1 | 0.987±0.021 | 0.451±0.091 |
|  | ESM-2_t12 | 33,563,302 | 375 | 0.976±0.025 | 0.473±0.055 |
| NA | WaveSeekerNet (Baseline) | 1,144,454 | 10 | 0.999±0.0 | 0.98±0.035 |
|  | WaveSeekerNet (FFT-only) | 837,827 | 7 | 0.999±0.0 | 0.968±0.056 |
|  | WaveSeekerNet (No KAN) | 541,022 | 9.5 | 0.999±0.0 | 0.887±0.161 |
|  | Transformer-FNET-64 | 19,275 | 0.56 | 0.97±0.021 | 0.249±0.045 |
|  | Transformer-FNET-128 | 52,427 | 0.63 | 0.975±0.019 | 0.236±0.064 |
|  | Transformer-MHA-64 | 291,851 | 0.82 | 0.965±0.018 | 0.345±0.138 |
|  | Transformer-MHA-128 | 611,915 | 0.9 | 0.988±0.011 | 0.392±0.154 |
|  | ESM-2_t6 | 7,541,845 | 82.3 | 0.997±0.007 | 0.674±0.131 |
|  | ESM-2_t12 | 33,563,085 | 175 | 0.977±0.044 | 0.622±0.113 |

### 5.3 WaveSeekerNet is robust to different up-sampling and down-sampling strategies

Our results showed that the choice of resampling approach has little impact on WaveSeekerNet’s ability to generalize in both the subtype and host source prediction tasks. Supplementary Tables S9-12 provide performance and statistical comparison of WaveSeekerNet with various hyperparameters, contrasting two training data resampling strategies. When predicting subtypes and host source using the high-quality independent datasets, the probability that the same type of model will have a mean F1-score (Macro Average) falling within the ROPE is often greater than 90% in the subtype prediction task (Supplementary Tables S9, S10) and usually over 50% in the host prediction task (Supplementary Tables S11, S12). For example, when predicting host source using the high-quality RNA sequences of the HA segment, the baseline model achieved an F1-score (Macro Average) of 0.969±0.008 if up-sampling and down-sampling were performed before cross-validation, versus 0.965±0.015 if they were only performed on each training split (Supplementary Table S11). These two scenarios had a 100% probability of being within the ROPE, indicating that the difference between the two resampling methods is negligible. Similarly, when tested on the high-quality RNA sequences, the baseline model achieved an F1-score (Macro Average) of 1.0±0.0 for both HA and NA subtype prediction in both resampling approaches (Supplementary Table S9).

However, generalization performance appeared more likely to differ when the low-quality independent datasets were used. In these tests, the probability of a difference between the up-sampling and down-sampling scenarios tends to be higher (over 50%). To assess for a directionality to this difference (e.g., if resampling is preferred before cross-validation), we conducted the trinomial test using the mean difference in the F1-score (Macro Average) of each scenario for each experiment conducted (e.g., HA subtype prediction using the RNA or protein sequences) [86,87]. After correcting for multiple comparisons using the Benjamini-Hochberg correction, the test indicated no significant preference in where resampling was applied (*p-value* ≥ 0.05, Supplementary Table S13). Finally, we analyzed the F1-scores for specific hosts and subtypes to assess for differences between strategies. Once again, the trinomial test indicated no significant preference (*p-value* ≥ 0.05, Supplementary Table S14), whether using a ROPE of 0.025 or the stricter ROPE of 0.01.

### 5.4 WaveSeekerNet can predict Influenza HA and NA subtypes with high accuracy

We compared WaveSeekerNet’s performance in predicting HA subtypes with the Transformer-only models, the pre-trained ESM-2 models, VADR, and BLASTp (Supplementary Figure S4 and Figure 3). When tested on the high-quality datasets of RNA and protein sequences, WaveSeekerNet achieved a minimum score of 0.97 across all evaluation metrics and hyperparameters (Supplementary Figures S4a, S4c), except when the Lookahead hyperparameter was not used. Disabling the Lookahead optimizer, the WaveSeekerNet only achieved an F1-score (Macro Average) of 0.872 on the high-quality dataset of RNA sequences. Notably, when WaveSeekerNet was trained using FCGR representation of RNA sequences (Figures 3a, 3b), it obtained F1-scores (Macro Average) of 1.0 and 0.998 for high-quality and low-quality datasets, respectively. In contrast, the Transformer-only models achieved maximum F1-scores (Macro Average) of 0.849 and 0.404 for high-quality and low-quality datasets of RNA sequences, respectively. Supplementary Table S15 reports the F1-scores for the specific HA subtypes, demonstrating WaveSeekerNet’s superiority in correctly identifying common and rare HA subtypes. For example, the H1 (*n_train_* = 56,593, *n_test_* = 19,599) and H8 (*n_train_* = 189, *n_test_* = 12) subtypes each achieved an F1-score of 1.0, contributing to WaveSeekerNet’s overall F1-score (Macro Average) of 1.0. When compared with sequence similarity search methods, WaveSeekerNet’s performance was comparable to VADR (Figures 3a, 3b) and BLASTp (Figures 3c, 3d). Notably, when tested on both high-quality and low-quality datasets of RNA sequences, VADR produced an F1-score (Macro Average) of 1.0. It is also notable that although the pre-trained ESM-2 models achieved high performance on the high-quality protein sequences (Figure 3c), WaveSeekerNet still significantly outperformed them when tested on the low-quality protein sequences (Figure 3d).

**Figure 3:**
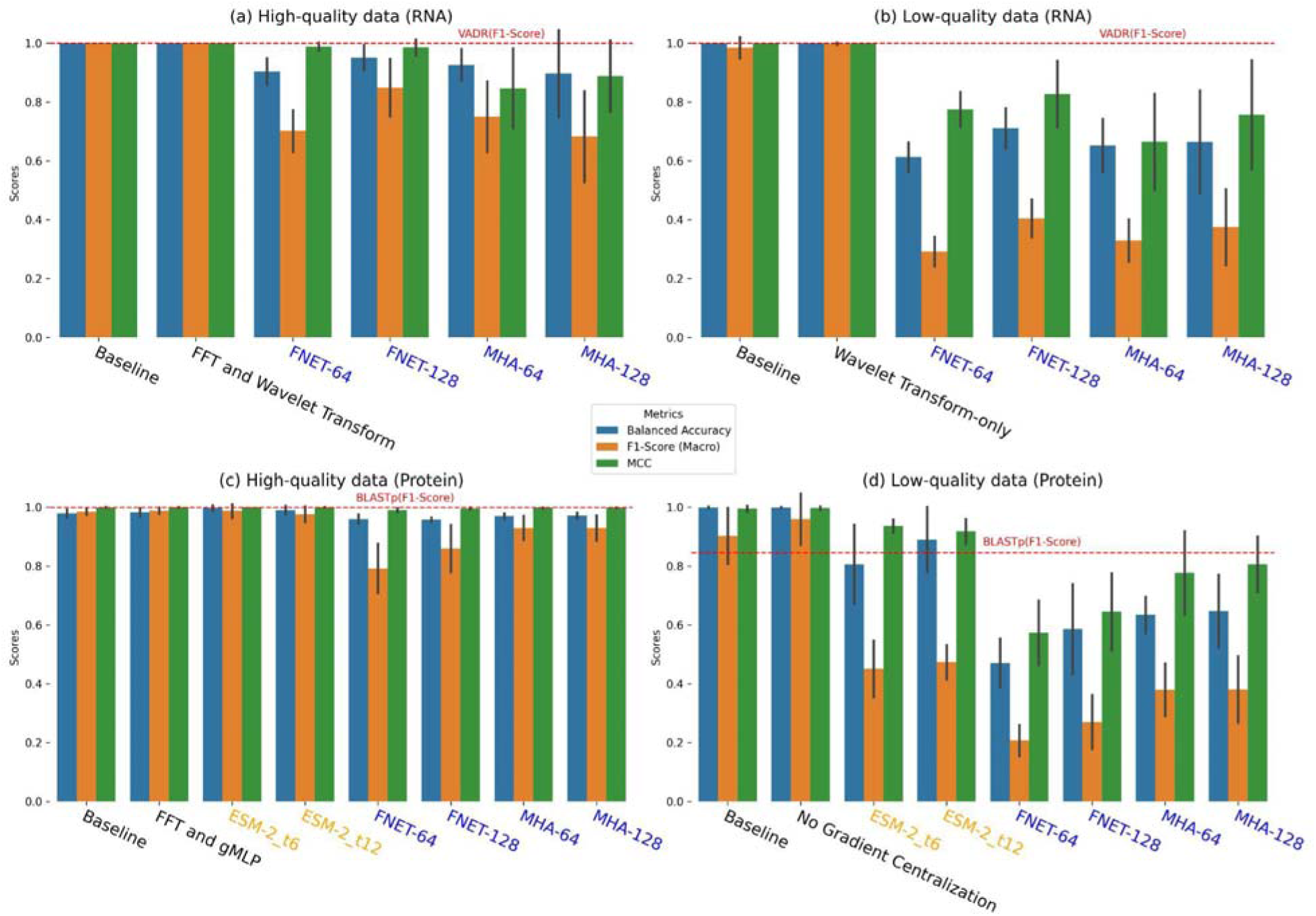
Performance comparison of HA subtype prediction when the best-performing WaveSeekerNet, pre-trained ESM-2, and Transformer-only models are tested. The performance of the baseline WaveSeekerNet is also shown as a point of reference. The bar plots of Balanced Accuracy, F1-score (Macro Average), and MCC are reported for the high-quality (a) and low-quality (b) FCGR representations of RNA sequences. The bar plots of scores for tests on the datasets constructed from high-quality and low-quality one-hot encoded protein sequences are reported in panels (c) and (d), respectively. Horizontal dashed lines present the F1-scores (Macro Average) for VADR and BLASTp. The WaveSeekerNet, pre-trained ESM-2, and Transformer-only models are labeled in Black, Orange, and Blue, respectively.

When classifying NA subtypes (Supplementary Figures S5, S6), the results mirrored the strong performance observed in HA subtype prediction. WaveSeekerNet achieved a minimum score of 0.995 across all evaluation metrics and hyperparameters on the high-quality datasets of RNA and protein sequences (Supplementary Figures S5a, S5c), except when the Lookahead hyperparameter was not used. Disabling the Lookahead optimizer, WaveSeekerNet only achieved an F1-score (Macro Average) of 0.818 on the high-quality dataset of RNA sequences (Supplementary Figure S5a). WaveSeekerNet also obtained an F1-score (Macro Average) of 1.0 on the high-quality dataset of RNA sequences, while the Transformer-only models achieved a maximum F1-score (Macro Average) of 0.966 (Supplementary Figure S6a). Supplementary Table S16 reports the F1-scores for the specific NA subtypes. Similarly to the HA subtype classification, both prevalent and rarer subtypes were accurately identified by WaveSeekerNet. For example, the F1-score for N2 (*n_train_* = 58,750, *n_test_* = 23,671) and N4 (*n_train_* = 347, *n_test_* = 41) was 1.0, contributing to WaveSeekerNet’s overall F1-score (Macro Average) of 1.0. When tested on the low-quality datasets of RNA and protein sequences, the Transformer-only models still performed the worst, with maximum F1-scores (Macro Average) of 0.728 and 0.392, respectively (Supplementary Figures S6b, S6d). In contrast, WaveSeekerNet maintained an F1-score (Macro Average) of 0.939, underperforming VADR, and a score of 0.998, outperforming BLASTp, when tested on the low-quality datasets of RNA and protein sequences, respectively. As expected, the performance of the dummy classifier was essentially random in each subtype prediction task. For example, using the high-quality RNA sequences, the dummy classifier achieved an F1-score (Macro Average) of 0.066 with NA subtype prediction.

### 5.5 WaveSeekerNet accurately identifies the host source using influenza A virus consensus sequences

Figures 4 and 5 show the comparisons between the best-performing WaveSeekerNet, pre-trained ESM-2, and Transformer-only models in predicting host source when tested on the HA segment and the combined HA and NA segments. We also evaluated the performance of the pre-trained ESM-2 model for host source prediction, using the protein sequences of the combined HA and NA segments (Figures 5c, 5d). Owing to the substantial GPU memory demands of the pre-trained ESM-2, particularly with this 2-channel configuration, the test was restricted to the ESM-2_t6 model, employing a training batch size of 32. When tested on the high-quality datasets of the HA segment, the best-performing WaveSeekerNet models achieved F1-scores (Macro Average) of 0.969 and 0.962 for RNA and protein sequences, respectively (Figures 4a, 4c). In contrast, the Transformer-only models achieved maximum F1-scores (Macro Average) of 0.917 and 0.916 for RNA and protein sequences, respectively. However, when evaluated on the high-quality datasets of the combined HA and NA segments (Figures 5a, 5c) and the NA segment (Supplementary Figures S7a, S7c), the best-performing Transformer-only models were comparable to WaveSeekerNet. The pre-trained ESM-2 models also achieved the same performance level as WaveSeekerNet for host source prediction using the high-quality protein sequences (Figures 4c, 5c, and Supplementary Figure S7c). The performance of the dummy classifier was essentially random, achieving an F1-score (Macro Average) of 0.267 when tested with the RNA sequences of the HA segment.

**Figure 4:**
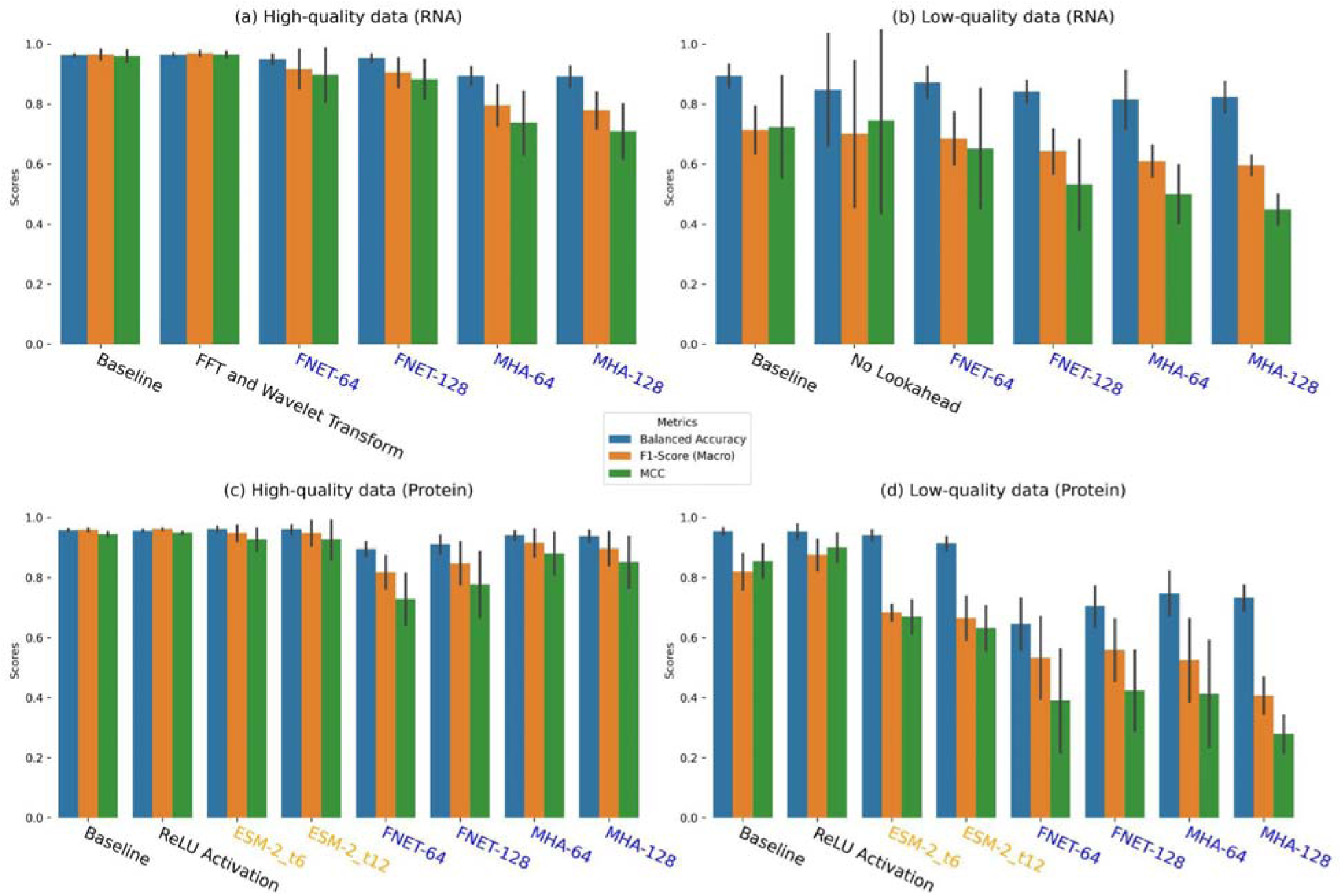
Performance comparison of host source prediction using the HA segment when the best-performing WaveSeekerNet, pre-trained ESM-2, and Transformer-only models are tested. The performance of the baseline WaveSeekerNet is also shown as a point of reference. The bar plots of Balanced Accuracy, F1-score (Macro Average), and MCC are reported for the high-quality (a) and low-quality (b) FCGR representations of RNA sequences. The bar plots of scores for tests on the datasets constructed from high-quality and low-quality one-hot encoded protein sequences are reported in panels (c) and (d), respectively. The WaveSeekerNet, pre-trained ESM-2, and Transformer-only models are labeled in Black, Orange, and Blue, respectively.

**Figure 5:**
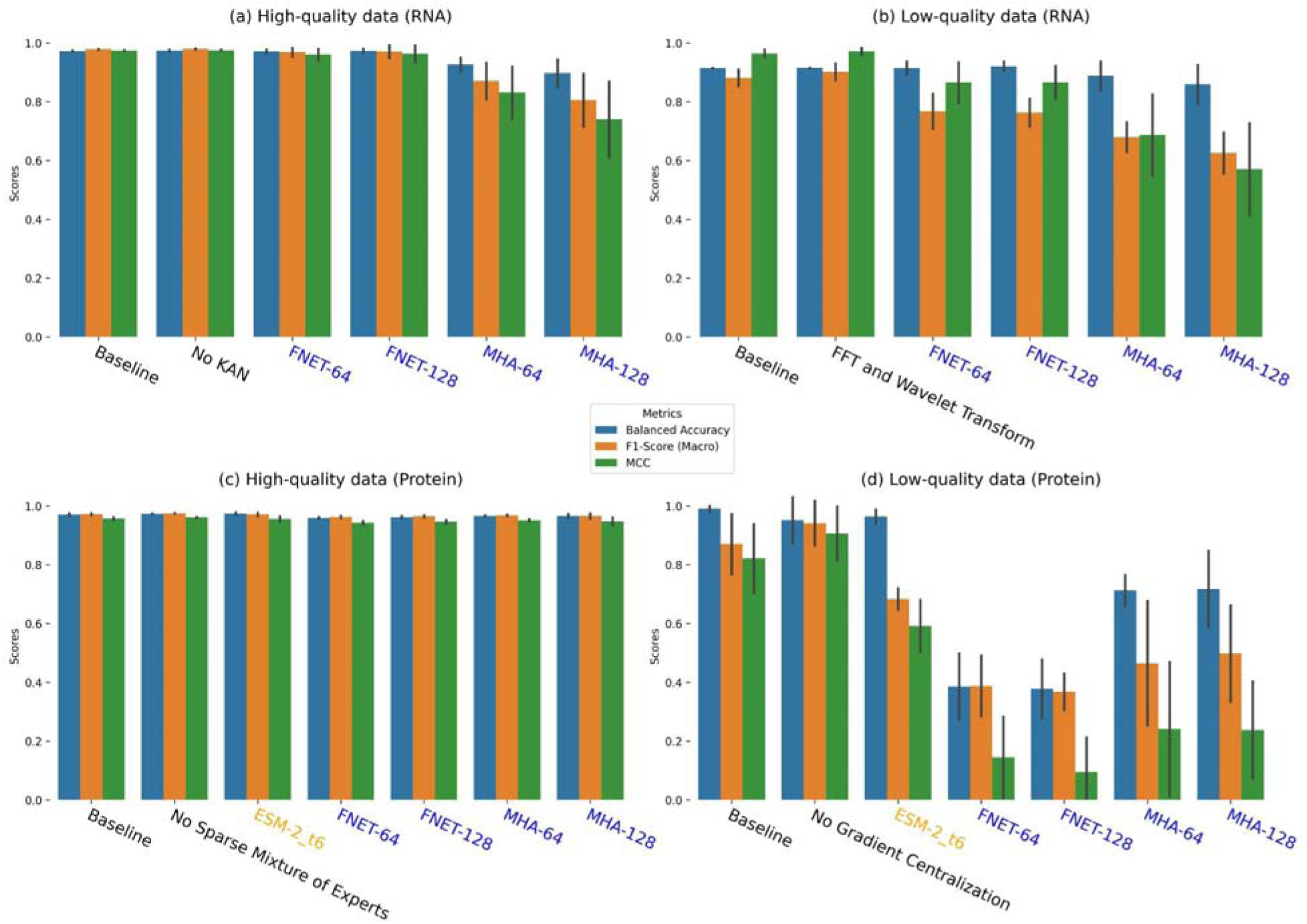
Performance comparison of host source prediction using the combined HA and NA segments (2 channels) when the best-performing WaveSeekerNet, pre-trained ESM-2, and Transformer-only models are tested. The performance of the baseline WaveSeekerNet is also shown as a point of reference. The bar plots of Balanced Accuracy, F1-score (Macro Average), and MCC are reported for the high-quality (a) and low-quality (b) FCGR representations of RNA sequences. The bar plots of scores for tests on the datasets constructed from high-quality and low-quality one-hot encoded protein sequences are reported in panels (c) and (d), respectively. The WaveSeekerNet, pre-trained ESM-2, and Transformer-only models are labeled in Black, Orange, and Blue, respectively.

A substantial difference in performance was observed when evaluating the low-quality datasets derived from protein sequences. WaveSeekerNet significantly surpassed the performance of the Transformer-only models by up to 44% in terms of F1-score (Macro Average) and by up to 67% in terms of MCC on the combined HA and NA segments (Figure 5d) and the HA segment (Figure 4d). The pre-trained ESM-2 models were significantly better than the Transformer-only models, but still lagged behind WaveSeekerNet (Figures 4d, 5d).

### 5.6 Host prediction discrepancies can carry important information about recent transmission events

In the previous sections, we demonstrated that WaveSeekerNet can accurately predict subtypes and host sources. Prediction discrepancies were noted and likely carry important information concerning recent transmission events since they imply that the extracted consensus sequence signature more closely resembles the signature from a different host. WaveSeekerNet produced multiple lines of evidence to support this hypothesis. For example, two H5N1 IAVs, *A/CastillaLaMancha/3739/2022* (EPI_ISL_15542438) and *A/CastillaLaMancha/3869/2022* (EPI_ISL_16813290), were isolated in humans but were classified as having an avian origin by WaveSeekerNet. Upon further investigation, we found that the Spanish Influenza National Reference Laboratory (NRL) linked these samples to an outbreak from a poultry farm where workers were likely infected by hens [88]. In another case, our approach identified the HA and NA genes from a sample *A/China/ZMD-22-2/2022* (EPI_ISL_15613648) as having an avian origin. This was the case since this strain is nested within clades of H3 and N8 genes isolated from ducks and other waterfowl [89]. In another instance, a sample isolated in March 2024, *A/Vietnam/KhanhhoaRV1-005/2024* (EPI_ISL_19031556), was correctly identified by WaveSeekerNet as having an avian origin. Contact tracing and viral characterization revealed that the 21-year-old man from Khanh Hoa Province, Vietnam, was exposed to wild birds and subsequently infected with an H5N1 avian influenza virus [90,91].

In addition, our results suggest that discrepancies can identify reverse zoonotic transmission events. For example, our model classified *A/swine/North Carolina/A02751330/2022* (EPI_ISL_16891306), *A/swine/Ohio/A02751292/2022* (EPI_ISL_16891307), *A/swine/Cambodia/PFC37/2021* (EPI_ISL_17885993), and *A/swine/Cambodia/PFC33/2020* (EPI_ISL_17886005) as human sequences. These discrepancies could be interpreted as these strains containing a genomic signature resembling other human sequences. Subsequent phylogenetic analyses and molecular characterization demonstrated that these sequences are closely related to those circulating in nearby human populations, suggesting a human-to-swine transmission event [92,93]. Supplementary Tables S17-19 provide additional details of discrepancies identified by our model.

### 5.7 WaveSeekerNet flags spillover events from the ongoing H5Nx avian influenza outbreak in North America

WaveSeekerNet successfully identified the most probable animal source of transmission in 1,659 sequences collected from the ongoing H5Nx avian influenza outbreak in North America. The model predicted that 100% of the sequences were of avian origin. For example, infections of mammals with the HPAI clade 2.3.4.4b H5N1, H5N5 viruses were recently detected [14,15,94]. These include H5N5 cases collected in Canada (e.g., *A/Raccoon/PEI/FAV-0199-1/2023* and *A/Striped_Skunk/PEI/FAV-0210-1/2023)*, H5N1 cases in dairy cows in the United States (e.g., *A/dairy_cow/Colorado/24_018028-008/2024*and *A/dairy_cow/Colorado/24_018028-013/2024*). Furthermore, a phylogenetic analysis supported the predictions for other H5N1 cases (e.g., humans, cats, dairy cows, goats, and swine). Thi analysis revealed that these samples are nested within avian sub-clades (Figure 6). Additionally, the collection sites of these samples are within the same geographical areas, providing additional evidence for a relationship between the avian strains and closely related strains isolated from non-avian hosts. Details of the sequences and predictions made by WaveSeekerNet are available in Supplementary Table S20.

**Figure 6:**
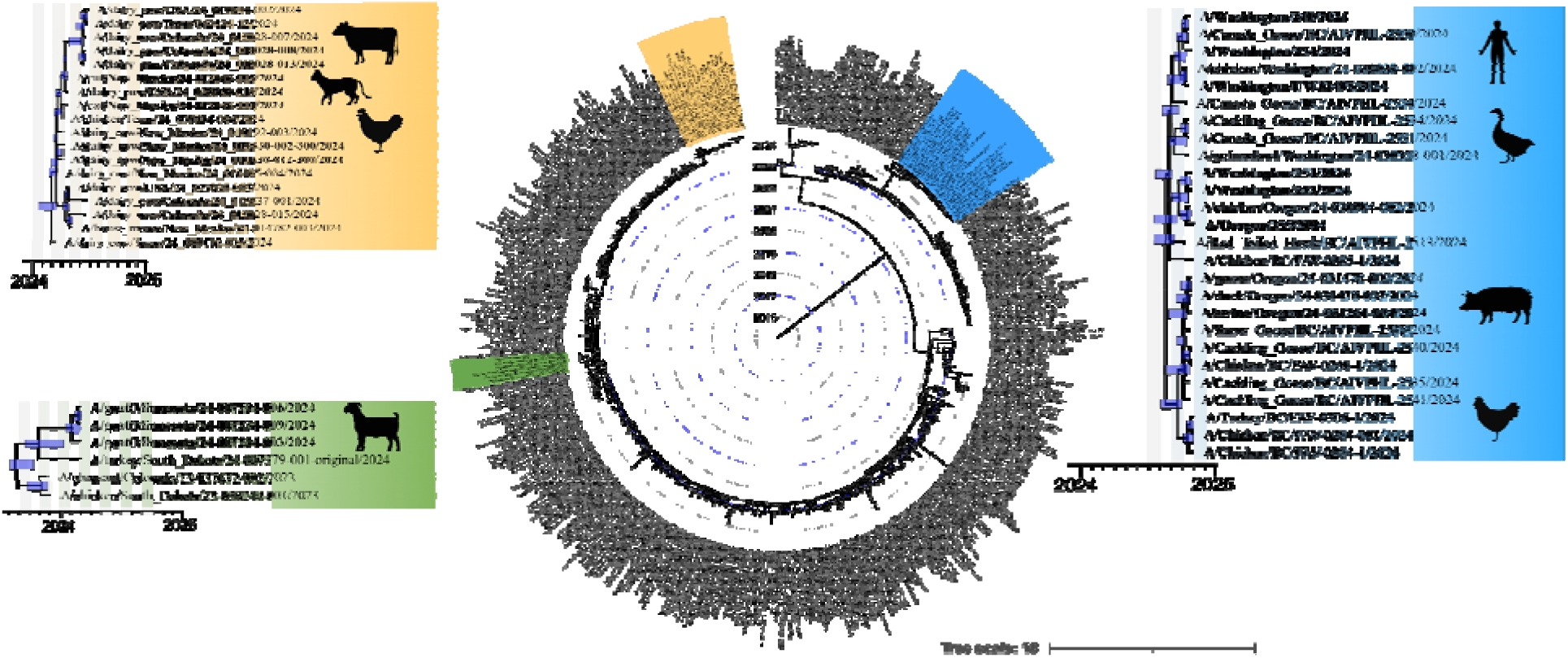
Pruned time-calibrated phylogeny of 1,659 HA RNA sequences collected from the ongoing H5Nx outbreak in North America. Different spillover events into mammalian hosts are highlighted in blue, orange, and green. WaveSeekerNet identified these strains as having an avian origin, a discrepancy supported by the phylogenetic analysis. Bars at the nodes indicate the 95% highest posterior density of the estimated node dates.

## 6 Discussion

This work presents a transformer-like architecture, WaveSeekerNet, inspired by ensemble learning, which can sufficiently capture and generalize information found in relatively simple sequence transformations and encodings, into a coherent internal representation of human, avian or non-human mammal-adapted IAV irrespective of subtype. The exceptional accuracy of WaveSeekerNet in predicting IAV subtypes and host source in both high-quality and low-quality test data underscores the role modern deep learning can play in rapid and effective influenza surveillance and characterization: especially with human activities such as hunting, agriculture, movement, and urbanization dramatically increasing interactions between humans and animal populations, subsequently increasing the risk of zoonotic transmission [95,96]. This increase in transmission risk is best exemplified by the emergence of the 2009 H1N1 IAV pandemic, which was a triple reassortant virus between human, avian, and swine strains [97], and the recent emergence of SARS-CoV-2 virus, which was associated with wildlife sold at the Huanan seafood market in Wuhan, China [98].

The approach presented here is advantageous since WaveSeekerNet was trained using simple sequence transformations and encodings. Simple transformations are effective since they are well studied and their applicability to problems such as these is known [25,38,39,46]. By utilizing these transformation methods, we can have greater confidence that any observed improvements can be attributed to our architectural choices rather than a lightly investigated and poorly understood choice of transformation. For example, the frequency of specific *k*-mers and codon pairs was found to be important for classifying potential hosts of H3Nx viruses [23]. Depending on the depth of the FCGR, these frequencies and features are natural features of the representation. Furthermore, by using the FCGR, we can also capture the unique fractal properties within the genome [43,99,100]. Together, this allows the model to learn from an information-dense set of features, which we demonstrate with our exceptional performance in subtype and host source prediction. Finally, since these transformations can be created by applying existing tools and do not require specialized knowledge, the workflow needed to run an analysis is greatly simplified. This is a critical requirement in diagnostic settings where the generation of accurate and reproducible reports for a timely and effective response to a disease outbreak is needed.

To test the efficacy of the different models, we used time-based data partitioning. This is a departure from the more traditional approach of random data partitioning that allows us to estimate the performance of a model better, since new viral strains are constantly emerging over time. Time-based data partitioning, therefore, can potentially minimize the likelihood of generating overly optimistic predictions since there is more novelty in the validation data due to the combined effect of antigenic drifts and shifts, exemplified by the emergence of novel reassortants with increased fitness in the H5Nx avian influenza outbreak in North America [14,101]. Generalizing when trained on imbalanced data is also important because recognizing rarer, emerging variants is essential for successful viral surveillance and characterization, which in turn informs public health responses. Often, this information is discarded, or analyses do not fully consider information from rarer variants and noisy data. This can potentially result in an overestimation of generalization performance. For example, Xu et al. considered rare subtypes (e.g., H15, H17, H18, N10, and N11) together as a single group when testing their Transformer-only models and used a weighted one-vs-all strategy to produce F1-scores [24,65]. However, this strategy is biased towards good performance in the majority classes and, as we show, can result in overly optimistic performance scores (Supplementary Tables S15, S16). Alternatively, up-sampling and down-sampling of under-represented and over-represented classes can be used to work with highly imbalanced data. Different biases can be introduced depending on where up-sampling and down-sampling occur, and the results can be overly optimistic towards minority classes [102]. In our experiments, WaveSeekerNet appears robust to different up-sampling and down-sampling strategies since the choice in strategy is unlikely to result in large differences in generalization performance. The ability to generalize patterns found in noisy, imbalanced data that evolves through time is vital in real-world scenarios since information is often missing. Approaches, such as those adopted in WaveSeekerNet, which are better at extracting key information from this type of data, can enhance the ability of public health authorities to detect and properly respond to outbreaks caused by emerging or rare strains.

WaveSeekerNet is an alternative to traditional tools used in bioinformatics, such as VADR. While robust, VADR’s performance is tied to the quality of underlying reference models and databases. Adapting VADR for novel subtypes typically requires highly trained experts since their input is necessary to update the reference models, databases, and underlying processes. WaveSeekerNet, on the other hand, learns to extract and use relevant features from the training data without expert guidance. The resulting model can then be used as part of an end-to-end process starting with consensus sequences and ending with viral subtypes and host source prediction. This is particularly advantageous when data is diverse and/or noisy. For example, our tests using low-quality datasets have demonstrated that it is possible to use deep learning to accurately predict viral subtypes and host source in diverse and noisy real-world datasets. This can save time and resources in analyzing degraded samples, which can be reallocated towards annotating and investigating truly interesting or problematic cases. While WaveSeekerNet requires retraining with new data so that novel subtypes are accounted for, it may be possible to streamline this process using recent advances in transfer learning and model fine-tuning [66,103,104].

WaveSeekerNet presents a compelling alternative to models that use the traditional self-attention mechanism. This advantage stems directly from WaveSeekerNet’s balancing of computational costs and excellent predictive performance. Depending on the choice of hyperparameters, WaveSeekerNet’s parameter count can be even less than that of the more commonly used multi-head self-attention-based model. This highlights another key advantage of WaveSeekerNet: it can be a scalable and flexible alternative for viral classification and host source prediction. Specifically, WaveSeekerNet can be trained using off-the-shelf consumer parts within a reasonable amount of time, meaning it can be used widely and, depending on the hyperparameter choices, without a high-performance cluster. More importantly, we show that a relatively simple approach, when compared to large pre-trained models such as ESM-2, can be efficiently used to develop classification models for medically relevant pathogens.

WaveSeekerNet’s ability to accurately classify host source, even when trained on diverse subtypes, suggests that the model is learning underlying genomic signatures linked to host adaptation and subtype. This is best exemplified by our model’s ability to accurately identify the correct subtype and host source in temporally separated data. Furthermore, we provide evidence that discrepancies between prediction and known host source likely indicate higher similarity with genomic signatures from another host. This could mean that viruses have had limited time to evolve within and adapt to a new host. For example, our phylogenetic analysis of IAVs associated with the ongoing H5Nx outbreak in North America reveals that the genomic signature found in sequences in non-human mammals is very similar to that of circulating avian strains, suggesting a recent introduction into the non-human mammal group. This observation is likely influenced by the fact that influenza viruses tend to accumulate mutations over time that optimize their fitness within a particular host species, leading to distinct genomic signatures associated with different hosts (e.g., avian, swine, humans) [105–107]. Therefore, if a virus jumps to a new host and retains a signature similar to its previous host, it suggests the jump was recent and there has not been sufficient time for the virus to undergo the extensive mutations needed to adapt fully to the new host environment.

WaveSeekerNet’s design is based on an ensemble attention-like mechanism that involves splitting viral sequences into ‘word’ patches and mixing them. This process allows the model to develop an internal representation of each class and leverages this representation to make predictions. Further work is needed to identify ‘word’ patches strongly impacting predictions, which can reveal important sequence properties underpinning viral adaptation and transmission in new hosts. This could be achieved by using Explainable AI (XAI) techniques, like SHAP (SHapley Additive exPlanations) [108], which measure how impactful a feature is on the output of a trained model. In this work, ‘word’ patches with high SHAP values can be interpreted as specific genetic determinants of host adaptation. Previously, SHAP values were used to find specific mutations in the genome of SARS-CoV-2 potentially associated with adaptation in deer and mink hosts [19]. While a similar approach can be taken here, additional care must be taken to understand how much each feature’s SHAP value changes with different initial parameterizations of WaveSeekerNet. Therefore, an optimal strategy that identifies maximally relevant features inextricably involves minimizing the impact of noise arising from differences in the initial state of WaveSeekerNet. Once found, the insights revealed using these tools could deepen our understanding of viral evolution and biology in novel hosts and provide a secondary, independent source of evidence to support and complement traditional phylogenetic analyses and experimentation using animal models.

To enhance the robustness and capacity of our model, we incorporated some new ideas that yielded promising results. For example, replacing the traditional feed-forward network within the WaveSeeker block with the MH-SMoE network led to a demonstrable improvement in the performance of WaveSeekerNet. This work also demonstrated the potential of the KAN network, a recently developed alternative to traditional feed-forward networks, in improving the identification of biological sequences in most tests. The applicability of KAN and MH-SMoE networks extends beyond the identification and characterization of IAV, and we believe that these approaches can be effectively applied to broader taxonomic identification problems, such as assigning taxonomic labels (e.g., species, genus, family) to DNA and RNA sequences generated by high-throughput sequencing technologies. While the baseline WaveSeekerNet model has exhibited strong performance, the impact of specific hyperparameters, such as the Wavelet Transform block, KAN, MH-SMoE, optimizer, and the choice of activation functions, highlights the importance of careful model design and hyperparameter tuning. Future work should focus on refining these hyperparameters and further exploring additional architectural enhancements to improve WaveSeekerNet’s accuracy and generalizability. Moreover, incorporating XAI techniques is essential to better understand the model’s decision-making processes, ultimately facilitating more effective model refinement and optimization.

WaveSeekerNet represents an advancement in applying deep learning for influenza virus classification and host source prediction. Its accuracy, efficiency, and potential for revealing novel biological insights position it as a valuable tool for future influenza surveillance and pandemic preparedness. We will integrate WaveSeekerNet into CFIA-NCFAD/nf-flu [109], an existing IAV analysis workflow at the Canadian Food Inspection Agency (CFIA) - National Centre for Foreign Animal Disease (NCFAD), which houses the World Organization for Animal Health reference laboratory for avian influenza. This integration will enable a more comprehensive analysis of IAV, ultimately contributing to enhanced Canadian and global IAV surveillance and pandemic preparedness. Furthermore, our work highlights the crucial role of deep learning in analyzing medically important data, such as genomic sequences and clinical records, derived from a wide spectrum of viral pathogens. This includes not only influenza virus, with its seasonal and pandemic potential, but also other viruses posing significant public health threats like rotavirus and SARS-CoV-2. The power of deep learning in this context stems from its ability to discover subtle, long-range patterns within complex and diverse datasets, leading to demonstrably improved prognostic accuracy in predicting disease severity, treatment response, or outbreak trajectories [110,111]. Given these capabilities, we foresee the broad application of WaveSeekerNet to other high-consequence viruses such as rotavirus and SARS-CoV-2, potentially aiding in rapid surveillance and outbreak prevention.

## 7 Limitations and Future Work

While this study presents a significant advancement in IAV prediction, there are limitations to address in future research:

- *Explainable AI*: Using explainable AI approaches would enhance the interpretability of WaveSeekerNet’s predictions, providing insights into the specific sequence features driving its classifications. This would also provide mutations associated with host prediction that could be verified experimentally.
- *Complex Transformations*: Exploring more complex input transformations, incorporating amino-acid properties, could further improve the model’s ability to capture subtle differences between strains and hosts.
- *Species-Level Breakdown*: Expanding host source prediction to a species-level breakdown within avian and non-human mammals would offer more granular insights for targeted surveillance and control measures.

By addressing these limitations, future iterations of WaveSeekerNet can provide even more powerful tools for understanding and combating IAV.

## Additional Files

**Figure S1:** The generalization performance of WaveSeekerNet for host source prediction was evaluated on the HA segment using various hyperparameter settings. The Balanced Accuracy, F1-score (Macro Average), and MCC are reported for high-quality (a) and low-quality (b) FCGR representations of RNA sequences. Scores for tests on the datasets constructed from high-quality and low-quality one-hot encoded protein sequences are reported in panels (c) and (d), respectively.

**Figure S2:** The generalization performance of WaveSeekerNet for host source prediction was evaluated on the NA segment using various hyperparameter settings. The Balanced Accuracy, F1-score (Macro Average), and MCC are reported for high-quality (a) and low-quality (b) FCGR representations of RNA sequences. Scores for tests on the datasets constructed from high-quality and low-quality one-hot encoded protein sequences are reported in panels (c) and (d), respectively.

**Figure S3:** The generalization performance of WaveSeekerNet for host source prediction was evaluated on the combined HA and NA segments (2 channels) using various hyperparameter settings. The Balanced Accuracy, F1-score (Macro Average), and MCC are reported for high-quality (a) and low-quality (b) FCGR representations of RNA sequences. Scores for tests on the datasets constructed from high-quality and low-quality one-hot encoded protein sequences are reported in panels (c) and (d), respectively.

**Figure S4:** The generalization performance of WaveSeekerNet and Transformer-only for HA subtype prediction using various hyperparameter settings. The Balanced Accuracy, F1-score (Macro Average), and MCC are reported for high-quality (a) and low-quality (b) FCGR representations of RNA sequences. Panels (c) and (d) show scores for high-quality and low-quality one-hot encoded protein sequences, respectively. The WaveSeekerNet and Transformer-only models using various hyperparameter settings are labeled in Black and Blue, respectively. The F1-scores (Macro Average) for VADR and BLASTp are shown as red horizontal dashed lines.

**Figure S5:** The generalization performance of WaveSeekerNet and Transformer-only for NA subtype prediction using various hyperparameter settings. The Balanced Accuracy, F1-score (Macro Average), and MCC are reported for high-quality (a) and low-quality (b) FCGR representations of RNA sequences. Panels (c) and (d) show scores for high-quality and low-quality one-hot encoded protein sequences, respectively. The WaveSeekerNet and Transformer-only models using various hyperparameter settings are labeled in Black and Blue, respectively. The F1-scores (Macro Average) for VADR and BLASTp are shown as red horizontal dashed lines.

**Figure S6:** Performance comparison of NA subtype prediction when the best-performing WaveSeekerNet, pre-trained ESM-2, and Transformer-only models are tested. The performance of the baseline WaveSeekerNet is also shown as a point of reference. The bar plots of Balanced Accuracy, F1-score (Macro Average), and MCC are reported for the high-quality (a) and low-quality (b) FCGR representations of RNA sequences. The bar plots of scores for tests on the datasets constructed from high-quality and low-quality one-hot encoded protein sequences are reported in panels (c) and (d), respectively. Horizontal dashed lines present the F1-scores (Macro Average) for VADR and BLASTp. The WaveSeekerNet, pre-trained ESM-2, and Transformer-only models are labeled in Black, Orange, and Blue, respectively.

**Figure S7:** Performance comparison of host source prediction using the NA segment when the best-performing WaveSeekerNet, pre-trained ESM-2, and Transformer-only models are tested. The performance of the baseline WaveSeekerNet is also shown as a point of reference. The bar plots of Balanced Accuracy, F1-score (Macro Average), and MCC are reported for the high-quality (a) and low-quality (b) FCGR representations of RNA sequences. The bar plots of scores for tests on the datasets constructed from high-quality and low-quality one-hot encoded protein sequences are reported in panels (c) and (d), respectively. The WaveSeekerNet, pre-trained ESM-2, and Transformer-only models are labeled in Black, Orange, and Blue, respectively.

**Table S1:** The data distribution of HA subtypes.

**Table S2:** The data distribution of HA sequences used for host source prediction.

**Table S3:** The data distribution of NA subtypes.

**Table S4:** The data distribution of NA sequences used for host source prediction.

**Table S5:** The data distribution of the combined HA and NA sequences used for host source prediction.

**Table S6:** The experimental results of host source prediction when testing the trained WaveSeekerNet using FCGR representation of RNA sequences.

**Table S7:** The experimental results of host source prediction when testing the trained WaveSeekerNet using one-hot encoding representation of protein sequences.

**Table S8:** Comparison of computational costs and efficiency for the WaveSeekerNet, Transformer-only, and pre-trained ESM-2 models when they were trained to predict host source using protein sequences of the HA and NA segments. For the WaveSeekerNet models, we show the baseline model with complete options and the modified baseline models using the fewest parameters.

**Table S9:** Performance and statistical comparison of WaveSeekerNet with various hyperparameters, contrasting two training data resampling strategies: (A) resampling training data within each cross-validation fold, (B) resampling the entire training data before cross-validation. The table provides F1-scores (Macro Average) for subtype prediction when testing the trained model using the high-quality and low-quality independent RNA sequences. The probabilities were computed using the Bayesian correlated t-test. A region of practical equivalence (ROPE) of 0.025 was used to quantify the probability that the overall generalization performance of models of each resampling approach differs by less than 0.025.

**Table S10:** Performance and statistical comparison of WaveSeekerNet with various hyperparameters, contrasting two training data resampling strategies: (A) resampling training data within each cross-validation fold, (B) resampling the entire training data before cross-validation. The table provides F1-scores (Macro Average) for subtype prediction when testing the trained model using the high-quality and low-quality independent protein sequences. The probabilities were computed using the Bayesian correlated t-test. A region of practical equivalence (ROPE) of 0.025 was used to quantify the probability that the overall generalization performance of models of each resampling approach differs by less than 0.025.

**Table S11:** Performance and statistical comparison of WaveSeekerNet with various hyperparameters, contrasting two training data resampling strategies: (A) resampling training data within each cross-validation fold, (B) resampling the entire training data before cross-validation. The table provides F1-scores (Macro Average) for host source prediction when testing the trained model using the high-quality and low-quality independent RNA sequences. The probabilities were computed using the Bayesian correlated t-test. A region of practical equivalence (ROPE) of 0.025 was used to quantify the probability that the overall generalization performance of models of each resampling approach differs by less than 0.025.

**Table S12:** Performance and statistical comparison of WaveSeekerNet with various hyperparameters, contrasting two training data resampling strategies: (A) resampling training data within each cross-validation fold, (B) resampling the entire training data before cross-validation. The table provides F1-scores (Macro Average) for host source prediction when testing the trained model using the high-quality and low-quality independent protein sequences. The probabilities were computed using the Bayesian correlated t-test. A region of practical equivalence (ROPE) of 0.025 was used to quantify the probability that the overall generalization performance of models of each resampling approach differs by less than 0.025.

**Table S13:** The two-tailed trinomial test results for each classification task using the high-quality and low-quality datasets. After applying the Benjamini-Hochberg correction for multiple comparisons, the tests indicated no significant preference regarding where resampling was applied.

**Table S14:** The two-tailed trinomial test results for each classification task for specific subtypes and hosts using the high-quality and low-quality datasets. Asterisks (*) indicate the absence of data for specific subtypes and hosts in the test sets. After applying the Benjamini-Hochberg correction for multiple comparisons, the tests indicated no significant preference regarding where resampling was applied.

**Table S15:** The report of HA subtype-specific performance when testing models using FCGR representation of RNA sequences. The F1-scores were obtained using the best-performing WaveSeekerNet and Transformer-only models. Models were trained to recognize the 18 HA subtypes (18 classes in the training data). Asterisks (*) indicate the absence of data for specific subtypes in the testing datasets. For these subtypes, we report an F1-score of N/A. F1-scores were calculated after 10 cross-validation folds and are presented for both high-quality and low-quality datasets.

**Table S16:** The report of NA subtype-specific performance when testing models using FCGR representation of RNA sequences. The F1-scores were obtained using the best-performing WaveSeekerNet and Transformer-only models. Models were trained to recognize the 11 NA subtypes (11 classes in the training data). Asterisks (*) indicate the absence of data for specific subtypes in the testing datasets. For these subtypes, we report an F1-score of N/A. F1-scores were calculated after 10 cross-validation folds and are presented for both high-quality and low-quality datasets.

**Table S17:** The report of host discrepancies identified by WaveSeekerNet when testing on the HA segment.

**Table S18:** The report of host discrepancies identified by WaveSeekerNet when testing on the NA segment.

**Table S19:** The report of host discrepancies identified by WaveSeekerNet when testing on the combined HA and NA segments.

**Table S20:** The report of host source identified by WaveSeekerNet when testing on 1,659 HA RNA sequences from the ongoing H5Nx outbreak in North America.

**Algorithm S1:** An algorithm to transform a DNA/RNA sequence into a chaos game representation.

**Algorithm S2:** Pseudo-code for the Fourier Transform block.

**Algorithm S3:** Pseudo-code for the Wavelet Transform block.

**Algorithm S4:** Calculation of the Router Z-loss.

**Algorithm S5:** Calculation of the KAN Regularization Loss.

## Disclosure of use of AI-assisted tools

In the revision phases, the authors used Google Gemini 2.5 Pro (preview) and Grammarly to improve the readability and check for misspellings and grammar mistakes. Prompts such as “check grammar” and “check grammar with academic writing” were used with Google Gemini 2.5 Pro (preview). The authors reviewed and edited the suggestions generated by tools and took full responsibility for the content of the publication.

## Data Availability

All additional supporting data are available in the GigaScience repository, GigaDB [112]. The RNA and protein sequences of the HA and NA segments of IAV can be downloaded from EpiFlu GISAID [35] after creating an account and accepting the terms of use. DOME-ML annotations are available in DOME registry [113].

## Availability of Source Code and Requirements

- Project Name: WaveSeekerNet: Accurate Prediction of Influenza A Virus Subtypes and Host Source Using Attention-Based Deep Learning
- Project home page: https://github.com/nhhaidee/WaveSeekerNet
- Software Heritage PID [114]: swh:1:snp:e8289fc61c0c4fc506502e80056bad26e80373a7
- RRID:SCR_026942
- biotools:waveseekernet
- Operating System(s): e.g, Platform independent
- Programming Language: Python 3.12.5
- Other requirements: Python 3.12+, pytorch 2.4.1, pytorch-optimizer 3.1.1, pytorch-wavelets 1.3.0, scikit-learn 1.5.1, seaborn 0.13.2, pyfastx 2.1.0, pandas 2.2.2, numpy 1.26.4, TrinomialTest 1.0.4
- License: MIT license

## Funding

H-H.N., J.R., C.L., O.V., and O.L. are supported by the Canadian Safety and Security Program (CSSP) grant CSSP-2022-CP-2538 (Artificial Intelligence for Front-Line Laboratories: Preparing for High-Consequence Pathogens to Reduce Their Risk). J.R., D.L., C.L., and O.L. are supported by the CSSP grant CSSP-2023-CP-2620 (Pilot Pan-Canadian Surveillance for Mammalian Viral Pathogens Using Hematophagic Organisms and Environmental Samples). H-H.N. and C.K.L. are partially supported by the Natural Sciences and Engineering Research Council of Canada (NSERC) and the University of Manitoba. G.W.T. and N.L. are supported by the INSPIRE (Integrated Network for the Surveillance of Pathogens: Increasing REsilience and capacity in Canada’s pandemic response) project funded through the Canada Biomedical Research Fund (grant CBRF2-2023-00008), the Biomedical Research Infrastructure Fund, and the Ontario Research Fund. This research was supported, in part, by the Province of Ontario and the Government of Canada through the Canadian Institute for Advanced Research (CIFAR), and companies sponsoring the Vector Institute (https://vectorinstitute.ai/partnerships/current-partners/).

## Author’s Contributions

H-H.N. designed the models, wrote the code/manuscript, prepared data, trained models, performed experiments and completed the data analysis. J.R. designed models, wrote the code, reviewed/edited the manuscript, mentored and provided critical feedback. O.V. performed phylogenetic analysis and reviewed/edited the manuscript. N.L. demonstrated the viability of the FCGR. O.L., C.K.L., and G.W.T. supervised the study and reviewed/edited the manuscript. D.L. and C.L. reviewed/edited the manuscript. All authors contributed to finalizing the manuscript.

## Acknowledgements

We thank Peter Kruczkiewicz, Cass Erdelyan, Dr. Anthony Signore, and Dr. Yohannes Berhane at the Canadian Food Inspection Agency (CFIA) - National Centre for Foreign Animal Disease (NCFAD) and reviewers for their valuable comments and feedback.

## Competing Interests

The authors have declared that no competing interests exist.

## Notes

https://gigadb.org/dataset/102732

### Competing Interest Statement

The authors have declared no competing interest.

### Summary of Updates

Updated to reflect comments by reviewers.

